# Organisation of afferents along the anterior-posterior and medial-lateral axes of the rat orbitofrontal cortex

**DOI:** 10.1101/2020.08.28.272591

**Authors:** Ines V. Barreiros, Marios C. Panayi, Mark E. Walton

**Affiliations:** Department of Experimental Psychology, University of Oxford, Tinsley Building, Mansfield Road, Oxford OX1 3SR, UK; Nuffield Department of Clinical Neurosciences, University of Oxford, John Radcliffe Hospital, Oxford OX3 9DU, UK; Wellcome Centre for Integrative Neuroimaging, University of Oxford, John Radcliffe Hospital, Oxford OX3 9DU, UK

**Keywords:** amygdala, neuroanatomy, rostral-caudal, rodent, submedius nucleus, thalamus

## Abstract

The orbitofrontal cortex (OFC) has been anatomically divided into a number of subregions along its medial-lateral axis, which behavioural research suggests have distinct functions. Recently, evidence has emerged suggesting functional diversity is also present along the anterior-posterior axis of the rodent OFC. However, the patterns of anatomical connections that underlie these differences have not been well characterised. Here, we use the retrograde tracer cholera toxin subunit B to simultaneously label the projections into the anterior lateral (ALO), posterior lateral (PLO), and posterior ventral (PVO) portions of the rat OFC. Our methodological approach allowed us to simultaneously compare the density and input patterns into these OFC subdivisions. We observed distinct and topographically organised projection patterns into ALO, PLO, and PVO from the mediodorsal and the submedius nuclei of the thalamus. We also observed different levels of connectivity strength into these OFC subdivisions from the amygdala, motor cortex, sensory cortices and medial prefrontal cortical structures, including medial OFC, infralimbic and prelimbic cortices. Interestingly, while labelling in some of these input regions revealed only a gradient in connectivity strength, other regions seem to project almost exclusively to specific OFC subdivisions. Moreover, differences in input patterns between ALO and PLO were as pronounced as those between PLO and PVO. Together, our results support the existence of distinct anatomical circuits within lateral OFC along its anterior-posterior axis.

## Introduction

Over recent years there has been an increasing understanding that the rodent orbitofrontal cortex (OFC) is not functionally homogeneous, particularly along its medial-lateral axis (see Barreiros et al., 2020; Bradfield & Hart, 2020; Izquierdo, 2017 for reviews). A recent review (Izquierdo, 2017) examined the relationship between the functions reported in rat OFC studies and the anatomical placement of the recording or manipulation sites. This revealed that functional heterogeneity can be mapped, to a certain degree, onto the divisions established by classical OFC parcellation methods. These classical parcellation studies define OFC subregions predominantly along the medial-lateral axis, including medial (MO), ventral (VO), ventrolateral (VLO), lateral (LO), dorsolateral (DLO) and agranular insular (AI) portions (Krettek & Price, 1977b; Price, 2006; Ray & Price, 1992).

While cytoarchitectural, neuroanatomical, and behavioural studies have mostly focused on the medial-lateral axis, recent reports suggest that there may also be important distinctions along the anterior-posterior axis. For example, Panayi & Killcross (2018) found that, while either anterior or posterior LO lesions impaired Pavlovian outcome devaluation, only posterior, but not anterior, LO lesions disrupted Pavlovian reversal learning. Another study revealed that the anterior but not posterior portion of MO is critical for inferring unobservable action-dependent outcomes and for behavioural response adaptation in outcome-devaluation tasks (Bradfield et al., 2018). These findings suggest that, rather than functionally uniform, the currently recognised OFC subregions might be composed of smaller structural and functional regions along its anterior-posterior axis. However, given the relative lack of clear cytoarchitectonic differences (Van De Werd & Uylings, 2008), it remains an open question whether there are anatomical distinctions that might underpin these functional differences.

Classically, the boundaries between prefrontal cortical regions, including OFC, have been defined by their specific projections patterns with the mediodorsal (MD) thalamus (Rose & Woolsey, 1948). Surprisingly, in comparison to other prefrontal cortical regions, there have been relatively few studies on the anatomical connectivity of the rat OFC, particularly looking at anterior versus posterior differences. Moreover, because there have been few studies systematically characterising the connectivity of different OFC subregions within the same subjects, inferences between subjects have often been necessary to make these comparisons.

Here, we test whether functional heterogeneity across the anterior-posterior axis of the OFC is underpinned by differences in anatomical connectivity, and with a focus on LO. Specifically, we test the hypothesis that there are distinct anatomical projections to the anterior lateral (ALO) and posterior lateral OFC (PLO) that underlie the functional dissociation reported by Panayi & Killcross (2018). We contrast these regions with the posterior ventral OFC (PVO) an anatomically adjacent portion of the OFC which is thought to be functionally distinct from LO (Balleine et al., 2011; Corwin et al., 1994). Specifically, we use the retrograde tracer cholera toxin subunit B (CTB) to simultaneously characterise the afferent projections to these regions. This approach allows us to establish whether any differences reflect (1) a gradient of afferent projections that may be organised topographically, or (2) a unique pattern of inputs that projects exclusively to a single subregion. We found that ALO and PLO receive both unique and topographically distinct gradients of cortical and thalamic inputs. These anterior-posterior differences within LO were as strong as the medial-lateral differences between PLO and PVO. Strikingly, robust and topographically distinct projections from the submedius thalamus were a key characteristic of all OFC subregions investigated. Overall, these anterior-posterior anatomical distinctions within lateral OFC support the emerging functional heterogeneity within this region.

## Experimental Procedures

### Subjects

A total of 14 male Lister Hooded rats (Envigo, UK), ~275-300 g in weight at the start of experiment (8-12 weeks old), were used in this study. Animals were housed in groups of 3 in polycarbonate cages, with ad libitum access to water and food. The housing room was maintained at a room temperature of 23 ± 1°C, humidity of 40 ± 10% and on 12-hour light/dark cycles beginning at 7 A.M., with lights during the day. All animal experimental procedures were approved and carried out in accordance with the British Home Office regulations and under the Animals (Scientific Procedures) Act 1986 (UK).

### Surgery

Animals were placed in vaporization chambers and anesthetised initially with 4% isoflurane (2 L/min O_2_) and maintained on 1–2% isoflurane (2 L/min O2) for the rest of the procedure. After induction, the head was shaved, and the rat was secured in a stereotaxic frame (Kopf Instruments). Body temperature was maintained at 37 ± 0.5°C with the use of a homoeothermic heating blanket. Corneal dehydration was prevented with application of ophthalmic ointment (Lacri-Lube, Allergan). The scalp was cleaned with diluted hibiscrub and 70% ethanol, and a local anaesthetic, bupivacaine (2 mg/kg), was applied to the area. The skin on top of the head was retracted and holes were then drilled in the skull for the injections. 50 nl of retrograde tracer CTB coupled to Alexa Fluor 488, 555 or 648 conjugates (C34775, C34776, C34778, Life Technologies) was microinjected through micropipettes with ~10-μm Ø pulled from thinwalled borosilicate glass capillaries (outer Ø: 1.14 mm, inner Ø: 0.53 mm; 3-000-203-G/X, Drummond Scientific) using a vertical micropipette puller (PE-2, Narishige) via an automated injector (Nanoject III, Drummond Scientific). Tracers were injected unilaterally, into the right hemisphere, at a rate of 1 nl/s. Each animal received injection of one of the 3 tracers into ALO, PLO, and PVO. Tracer-site combinations were counterbalanced across animals. Coordinates used were as indicated in Table 1. Before injection, the micropipette was lowered 0.2 mm below the intended injection sites and left in place for 1 min, before being raised to the injection site dorsoventral coordinate. Micropipettes were left in place for 5 min after each bolus injection to ensure diffusion of tracer. Micropipettes were then slowly retracted, and the incision closed with vicryl sutures.

**Table 1.**
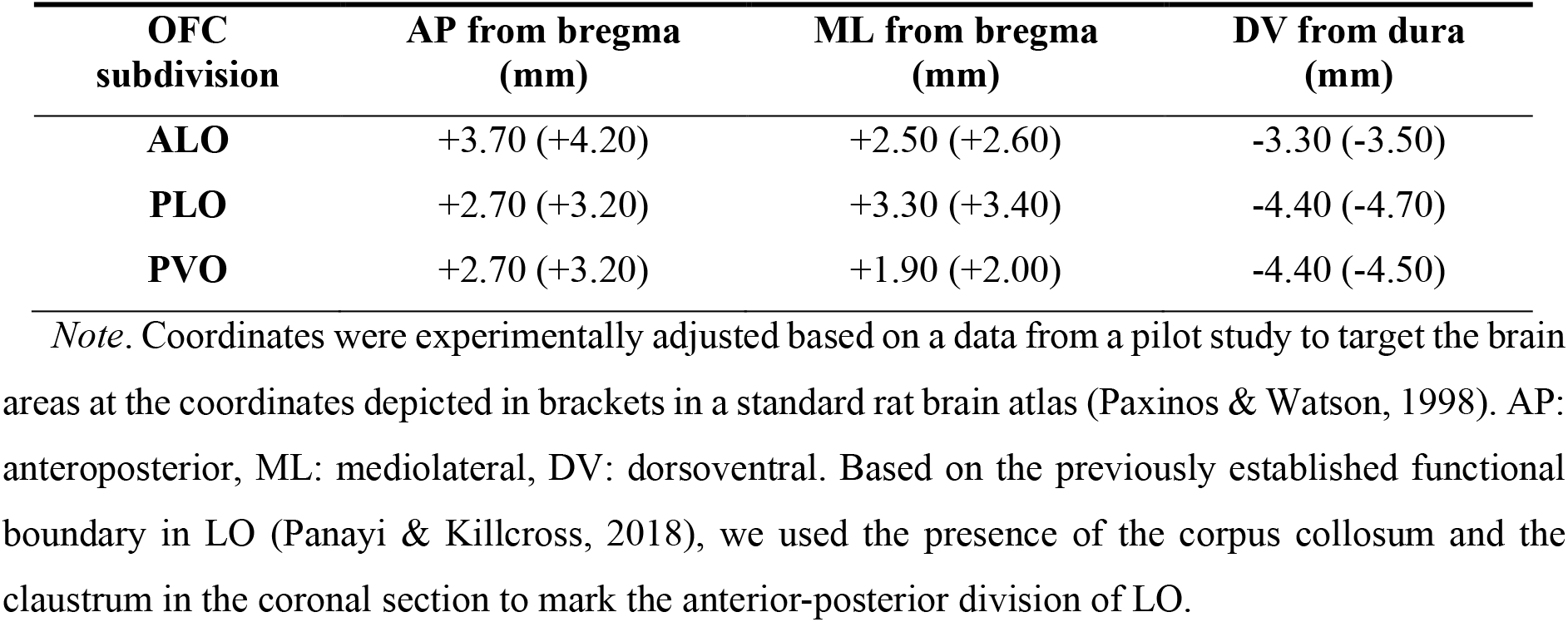
Coordinates used for tracer injection into OFC subdivisions

| OFC subdivision | AP from bregma (mm) | ML from bregma (mm) | DV from dura (mm) |
| --- | --- | --- | --- |
| **ALO** | +3.70 (+4.20) | +2.50 (+2.60) | -3.30 (-3.50) |
| **PLO** | +2.70 (+3.20) | +3.30 (+3.40) | -4.40 (-4.70) |
| **PVO** | +2.70 (+3.20) | +1.90 (+2.00) | -4.40 (-4.50) |

*Note.* Coordinates were experimentally adjusted based on a data from a pilot study to target the brain areas at the coordinates depicted in brackets in a standard rat brain atlas (Paxinos & Watson, 1998). AP: anteroposterior, ML: mediolateral, DV: dorsoventral. Based on the previously established functional boundary in LO (Panayi & Killcross, 2018), we used the presence of the corpus collosum and the claustrum in the coronal section to mark the anterior-posterior division of LO.

All animals were administered buprenorphine (0.1 ml/kg, s. c.), both pre- and post-surgically, and meloxicam (Metacam, 0.2/ml/kg, s. c.), post-surgically. Animals were then allowed to recover in thermostatically controlled cages and given palatable food for consumption. Meloxicam was also administered for at least 3 days following surgery.

### Histology

Ten days after tracer injection, animals were administered a terminal dose of pentobarbitone (30-60 mg/kg). After loss of pedal reflex, the animals were perfused transcardially with 150 ml of phosphate buffer saline followed by 400 ml of 4% paraformaldehyde. The brains were removed, kept in paraformaldehyde for 24 h and then transferred to phosphate buffer saline. Before slicing, brains were cryoprotected in 30% sucrose solution. 60-μm thick coronal sections were obtained with a freezing microtome (Leitz). Sections were mounted onto slides and coverslipped with mounting medium with DAPI (H-1500 Vectashield, Vector Labs).

### Image acquisition and analysis

Images of whole brains sections were captured using a fluorescence slide scanner (Axio Scan.Z1, Zeiss) equipped with an air 20x/NA 0.8 objective. Images of example single-, double, and triple-labelled neurons in the submedius nucleus of the thalamus were acquired using a confocal microscope (Zeiss LSM 880) equipped with a 20x/NA 0.8 Plan-Apochromat objective. Image colour settings, including brightness, contrast and gamma, were adjusted to aid visualisation of labelled cells. Levels of cell labelling density were manually categorised, offline, into four levels: 0, absence of labelling; 1, weak (i.e. just a few labelled cells); 2, moderate; 3, strong (see Figure S2 for examples). The summary table and figure depicting the average labelling for each injection site are based on the density average across brains, subsequently averaged across sections when a single section is represented and discretised into four density values: –, absence of labelling; +, weak; ++, moderate; +++, strong (Table 2 and Figure 8). The weak density level (+) also includes any region that is consistently labelled, i.e. across at least 50% of the brains, even if only weak labelling was observed. Areas with maximal tracer deposit, only found around injection sites, were not included in the analyses. If a brain region contained maximal tracer deposit in a coronal section in more than 50% of the brains included in the analyses then we represented that area as maximal deposit in the average labelling figure, otherwise we calculated the average labelling for that region excluding those brains. The classification of brain areas was based on a standard rat brain atlas (Paxinos & Watson, 1998). The nomenclature used to classify area 1 of the anterior cingulate cortex was based on (Paxinos & Watson, 2013), to reflect current understanding of the heterogeneity within the medial prefrontal cortex (Laubach et al., 2018). In our analysis, we focused on the quantification of frontal lobe structures, including OFC subdivisions, prelimbic, infralimbic, anterior cingulate cortex and motor cortex; temporal lobe structures, including amygdala, lateral entorhinal cortex and perirhinal cortex; retrosplenial cortex; primary sensory cortices; thalamic nuclei, including paratenial, submedius and mediodorsal. Due to difficulties in obtaining brain sections of consistent quality posterior to −5.30 mm from bregma, we did not quantify labelling in midbrain structures.

**Table 2.** Density of labelling in key areas observed following CTB injection into ALO, PLO, or PVO

| Structures | ALO | PLO | PVO |
| --- | --- | --- | --- |
| **Prefrontal cortex** |  |  |  |
| orbitofrontal cortex, medial part | - | + | + |
| prelimbic cortex | - | + | + |
| infralimbic cortex | - | + | + |
| anterior cingulate cortex, area 1 | - | - | + |
| motor cortex | ++ | - | + |
| **Sensory cortices** |  |  |  |
| granular insular cortex | + | + | + |
| piriform cortex | ++ | ++ | ++ |
| primary somatosensory cortex |  |  |  |
| *barrel cortex* | + | - | ++ |
| *forelimb, hindlimb & trunk areas* | - | - | + |
| secondary visual cortex |  |  |  |
| *lateral area, mediolateral & mediomedial areas* | - | - | + |
| auditory cortex | - | - | - |
| **Thalamic nuclei** |  |  |  |
| mediodorsal |  |  |  |
| *medial part* | + | ++ | - |
| *central part* | + | ++ | + |
| *lateral part* | ++ | + | + |
| submedius |  |  |  |
| *anterior section (-2.12\*)* |  |  |  |
| *dorsal part* | + | - | ++ |
| *ventral part* | + | ++ | + |
| *middle section (-2.30\*)* |  |  |  |
| *dorsal part* | ++ | + | +++ |
| *ventral part* | ++ | +++ | +++ |
| *posterior section (-2.56\*)* |  |  |  |
| *dorsal part* | ++ | + | + |
| *ventral part* | + | ++ | + |
| paratenial | - | ++ | - |
| reuniens | - | + | + |
| **Amygdala** |  |  |  |
| basolateral |  |  |  |
| anterior part | + | ++ | + |
| posterior part | - | + | - |
| lateral | - | + | - |

*Note.* Only regions where main differences and relevant similarities were observed are represented. Because we observed no consistent differences in labelling density between the anterior and posterior portions of MO, here we collapsed its average labelling across all coronal sections. -, absence of labelling; +, weak labelling; ++, moderate labelling; +++, strong labelling; \*, distance from bregma in mm; CTB: cholera toxin subunit B.

The proportion of double- and triple-labelled cells in the submedius nucleus of the thalamus was obtained by counting the number of single-, double-, and triple-labelled cells in this region in the coronal sections at −2.12, −2.30 and −2.56 mm from bregma of successfully triple-labelled brains. These values were then summed across sections and averaged across brains.

In the description and interpretation of our results, we consider anterior and posterior divisions of MO, VO, and LO separately, but not of DLO. We refer to the portions of LO and VO contained in the coronal sections from +4.70 to + 4.20 mm from bregma as anterior and to those contained in the coronal sections from +3.20 to +2.20 mm as posterior, with the corpus collosum and the claustrum in the coronal section marking the anterior-posterior division as previously established for LO (Panayi & Killcross, 2018). Based on a previously observed functional dissociation in MO (Bradfield et al., 2018), we refer to the areas contained in coronal sections at +4.70 and +3.70 mm from bregma as anterior and posterior MO, respectively. These coordinates are based on a standard rat brain atlas (Paxinos & Watson, 1998). While an area VLO has been proposed to exist between VO and LO, here we will only differentiate between areas VO and LO as per the boundaries defined in Paxinos & Watson (1998).

We tested whether differences in labelling strength were statistically different in key areas of interest (i.e. thalamus, amygdala) using the average labelling density across brain sections for each area subregion. We used a non-parametric approach to an ANOVA model to capture the factorial structure of the data analysed using R statistical software (Lenth et al., 2020; R Core Team, 2020). A non-parametric approach was important given that the labelling was quantified with ordinal data. We used an aligned rank transform procedure (Wobbrock et al., 2011) that allows for valid main effect and interaction inferences while maintaining a nominal Type I error rate. Multiple-comparison post-hoc analyses were corrected using a Sidak correction. In all analyses a mixed effects model with a random effect of subject was defined at the level of each brain, and tests are reported with Kenward-Roger residual degrees of freedom. We used a similar approach to statistically test differences in labelling strength in the subregions of the submedius nucleus of the thalamus across coronal sections.

Depiction of labelling in key areas of interest, including the amygdala, mediodorsal thalamus and submedius nucleus of the thalamus, was obtained by superimposing hand drawings of the labelling in each brain using an opacity value proportional to the density level of labelling observed. Data from injections that were off target (*n* = 11 injection sites), in which tracer did not diffuse (*n* = 8 injection sites), were both off target and the tracer did not diffuse (*n* = 6 injection sites), or in which the tracer diffused into the white matter (*n* = 2 injection sites) were excluded from all analyses (see Figure S1). The data presented here was obtained from 7 brains: 4 triple-labelled (*n* = 12 injection sites), 3 single-labelled (*n* = 3 injection sites).

## Results

### Simultaneous characterisation of ALO, PLO, and PVO afferents

We set out to simultaneously characterise the inputs of three distinct portions of the rat OFC: ALO, PLO, and PVO. We compared the projections of the three subdivisions in the same brain, while using the same retrograde tracer (cholera toxin subunit B, CTB), coupled to different fluorescent dyes (Conte et al., 2009). Figure 1 illustrates the intended injection sites and the core of tracer deposits observed following histological analysis of injected brains. The location of the core injection deposits included in the analysis were mostly confined to the intended OFC subdivision, with some extension into the ventral agranular insula (AIv) from the PLO injections. The localisation and average density of retrogradely-labelled cells observed are illustrated in Figure 2 (see Table S1 in the supplementary material for exact average density values). Labelling in each individual brain following tracer deposition into ALO, PLO and PVO is represented in Figures S3, S4, and S5, respectively, in the supplementary material.

**Figure 1.**
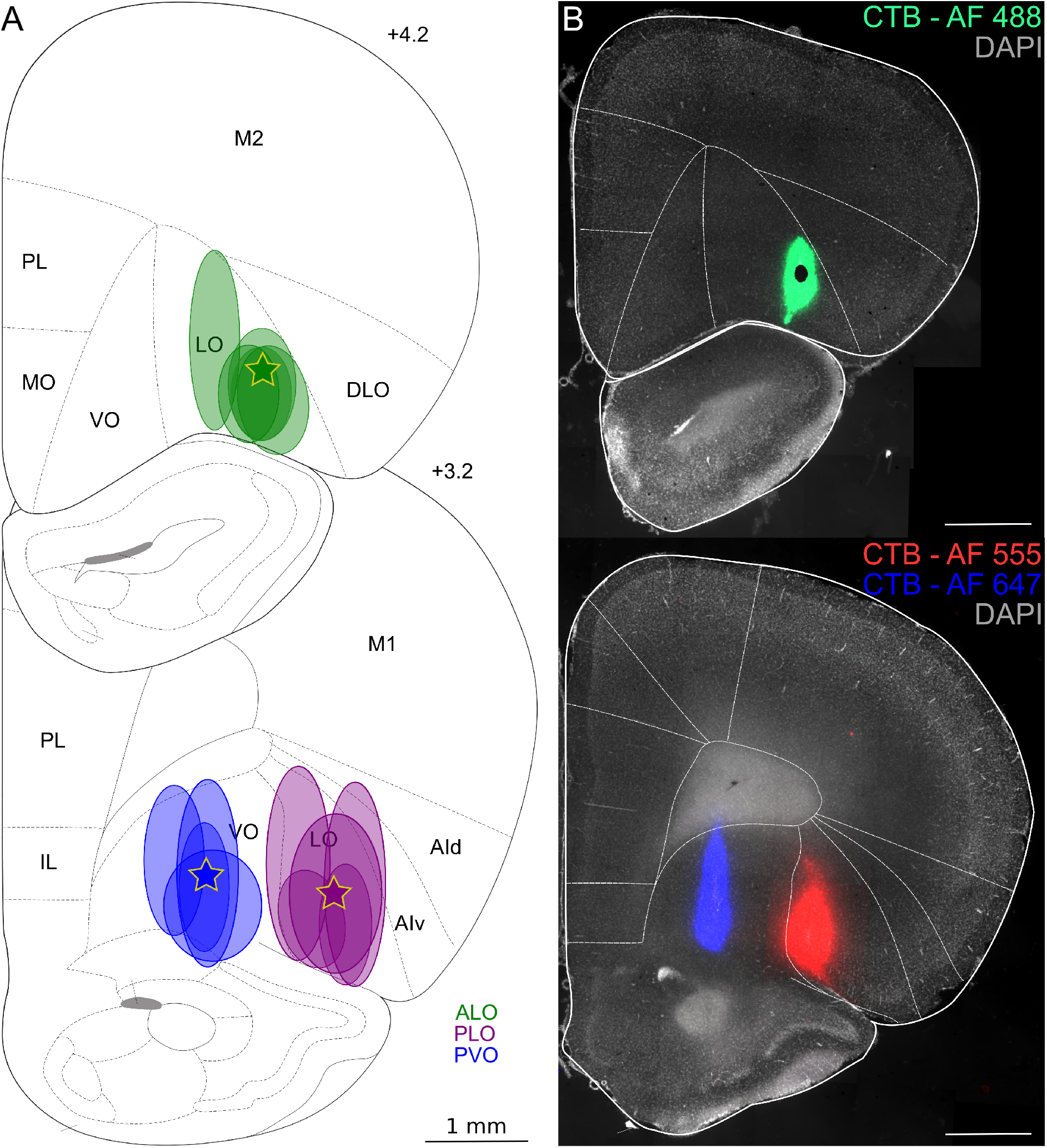
Coronal sections depicting core of tracer deposits into ALO, PLO, or PVO, illustrated at the level of maximal tracer deposit. (A) Stars and oval shapes represent the intended and the observed injection sites core, respectively. Injections whose core fell outside the orbitofrontal cortex or in which the tracer did not diffuse are not represented here (see instead Figure S1). *n* ALO = 6, *n* PLO = 5, *n* PVO = 4 injections. (B) Example injection sites of triple-labelled brain. Distances shown are distances from bregma in mm. AF: Alexa Fluor; AId: agranular insular cortex, dorsal part; AIv: agranular insular cortex, ventral part; ALO: lateral orbitofrontal cortex, anterior part; CTB: cholera toxin subunit B; DLO: orbitofrontal cortex, dorsolateral area; IL: infralimbic cortex; MO: orbitofrontal cortex, medial part; LO: orbitofrontal cortex, lateral part; M1: primary motor cortex; M2: secondary motor cortex; PL: prelimbic cortex; PLO: lateral orbitofrontal cortex, posterior part; PVO: ventral orbitofrontal cortex, posterior part; VO: orbitofrontal cortex, ventral part.

**Figure 2.**
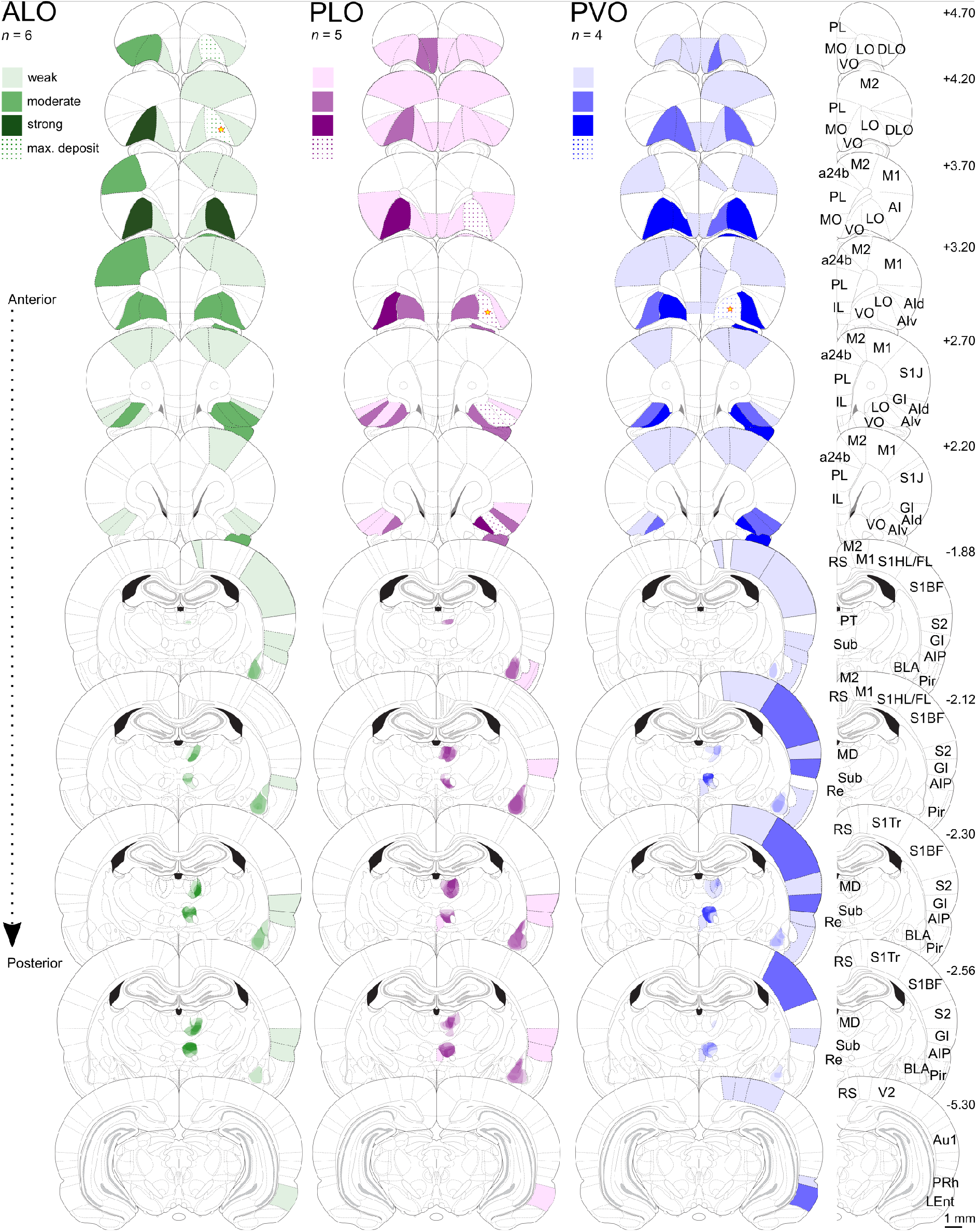
Density of retrogradely-labelled cells following CTB injection into ALO, PLO, or PVO. Represented is the average density across brains discretised into four labelling levels (see Table S1 for exact average density values). Lighter or darker shades represent lower or higher labelling densities, respectively. This represents the average density of projections from multiple brains within a given structure but note that this does not indicate the precise location of these projections within the structure. Depiction of labelling in key areas of interest (amygdala, mediodorsal thalamus and submedius nucleus of the thalamus) was obtained by superimposing hand drawings of the labelling in each brain using an opacity value proportional to the density level of labelling observed. Star shapes represent the intended injection site cores. Distances shown are distances from bregma in mm. *n* ALO = 6, *n* PLO = 5, *n* PVO = 4. a24b: anterior cingulate cortex, area 24b; ALO: lateral orbitofrontal cortex, anterior part; AI, v, d, P: agranular insular cortex, ventral area, dorsal area, posterior area; Au1: primary auditory cortex; BLA: basolateral amygdala; CTB: cholera toxin subunit B; DLO: dorsolateral orbitofrontal cortex; GI: granular insular cortex; IL: infralimbic cortex; LEnt: lateral entorhinal cortex; max. deposit: maximal deposit; LO: lateral orbitofrontal cortex; M1: primary motor cortex; M2: secondary motor cortex; MD: mediodorsal nucleus of the thalamus; MO: medial orbitofrontal cortex; Pir: piriform cortex; PL: prelimbic cortex; PLO: lateral orbitofrontal cortex, posterior part; PVO: ventral orbitofrontal cortex, posterior part; PRh: perirhinal cortex; PT: paratenial nucleus of the thalamus; Re: nucleus reuniens of the thalamus; RS: retrosplenial cortex; S1BF: primary somatosensory cortex, barrel field; S1J: primary somatosensory cortex, jaw region; S1HL/FL: primary somatosensory cortex, hindlimb and forelimb regions; S1Tr: primary somatosensory cortex, trunk region; S2: secondary somatosensory cortex; Sub: submedius nucleus of the thalamus; V2: secondary visual cortex; VO: ventral orbitofrontal cortex.

In our study, contralateral labelling was limited to prefrontal afferents (ending at +2.20 mm from bregma; Figure 2). The remaining labelling was observed only ipsilaterally. In the triple-labelled brains the vast majority of cells were single-labelled with only a small proportion of cells being double- or triple-labelled (see Figure 3 for examples). These were found most often in the submedius nucleus of the thalamus where we observed an average of 11% and 1% of labelled cells were double- and triple-labelled, respectively. The density of the retrogradely-labelled cells observed in key areas of interest is summarised in Table 2.

**Figure 3.**
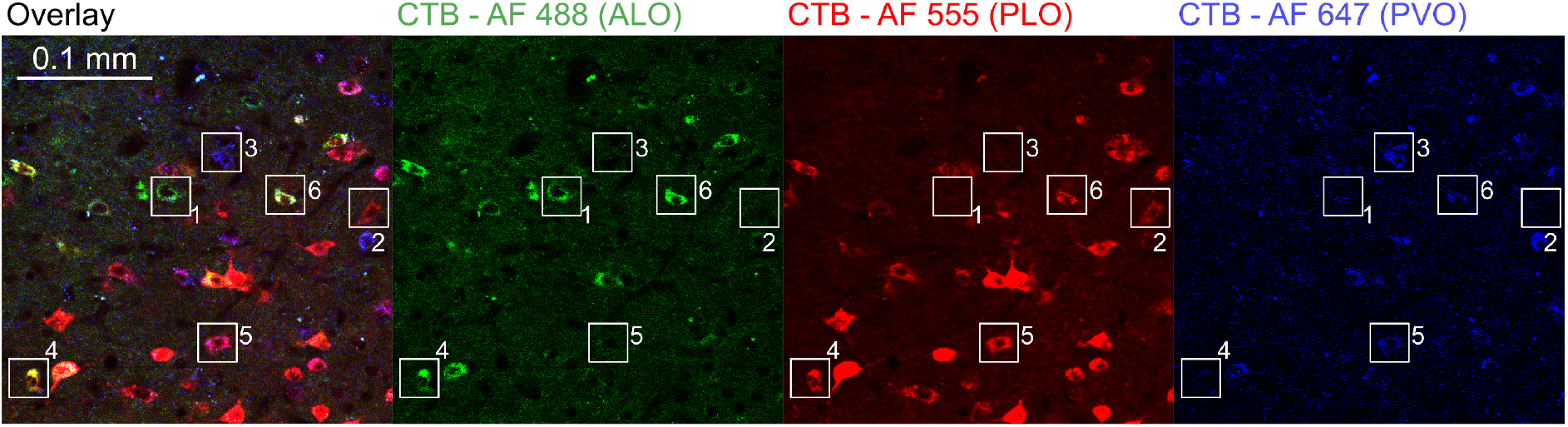
Example single-, double-, and triple-labelled cells. Represented is a plane of a z-stack obtained with a confocal microscope of the submedius nucleus of the thalamus in which are present: single-labelled cells after (1) ALO, (2) PLO, or (3) PVO injections; double-labelled cells after (4) ALO and PLO, or (5) after PLO and PVO injections; and (6) triple-labelled cells.

### Prefrontal cortex afferents

#### Intra-OFC projections

##### DLO

following injection of CTB into ALO and PLO, we observed weak to moderate labelling in DLO, ipsi- and contralaterally to the injection site (Figure 2). In contrast, we detected no labelling in this OFC portion following injection into PVO, both ipsi- and contralaterally.

##### AI

following ALO injection, we detected only weak labelling in the AI of the same coronal section as the injection site and weak to moderate labelling in posterior sections. PLO injection resulted in weak or strong ipsilateral labelling in anterior and posterior AI, respectively. Retrograde labelling in the contralateral side was mostly weaker. PVO injection resulted in weak to moderate density of labelled cells in the posterior portions of AI.

##### LO

ALO injections of CTB resulted in strong or moderate labelling in ALO and PLO, both ipsi- and contralaterally. PLO injections resulted in only weak labelling in ALO, ipsilaterally. Contralaterally, we detected weak to moderate labelling in ALO and strong to weak labelling in PLO, along the anterior-posterior axis. Following injection of CTB into PVO, we observed strong to moderate labelling in PLO and only weak to moderate labelling in ALO, ipsilaterally and along the anterior-posterior axis. This was largely mirrored on the contralateral hemisphere.

##### VO

Following tracer injection into ALO, we detected only weak labelling in anterior ventral OFC (AVO) but moderate labelling in PVO, ipsilaterally. A similar labelling pattern was observed following tracer injection into PLO. This was largely mirrored on the contralateral hemisphere both in ALO and PLO injections. Injection into PVO resulted in weak or strong labelling in ipsilateral AVO or PVO, respectively. Retrograde labelling in the contralateral side largely mirrored that of the ipsilateral cortex, generally with fewer cells present.

##### MO

MO had almost no labelled cells following ALO injection, with only some very low-density labelling in its most anterior part in both hemispheres in 2 of the 6 brains included in the analysis (see Figure S3). By contrast, both PLO and PVO injections resulted in weak but consistent labelling in both the anterior and posterior portions of MO, ipsilaterally. A similar pattern was observed in the contralateral hemisphere, with the exception that the anterior portion of MO contained stronger labelling contralateral to the injection site following PVO injection.

In summary, all three subdivisions receive input from most other parts of the OFC, both ipsi- and contralaterally. One distinction is MO, which sends stronger projections to PLO and PVO than to ALO. By contrast, DLO sends weak projections to both ALO and PLO but not to PVO.

#### Medial prefrontal projections

Following retrograde tracer injection into ALO, prelimbic (PL) and infralimbic (IL) were mostly devoid of labelled cells, both ipsi- and contralaterally (Figure S3). On the other hand, following PLO injection, both IL and PL contained weak labelling in a majority of brains. Similarly, we observed weak but consistent labelling in both IL and PL following PVO injection. No labelled cells were detected consistently in the anterior cingulate cortex (a24b), in either hemisphere following injection into ALO or PLO. By contrast, PVO injection often resulted in weak labelling in this region (Figure 2, Figure S5). Thus, IL and PL labelling suggest an anterior-posterior difference within LO, while a24b labelling suggests a medial-lateral distinction between PLO and PVO (Table 2).

#### Motor cortex

Following injections into ALO, we observed weak but widespread labelling in both ipsilateral, primary (M1) and secondary motor (M2) cortex (Figure 2). This was largely mirrored in the contralateral side with the difference that, here, ALO resulted in moderate labelling in M1. By contrast, PLO injection resulted only in weak labelling restricted to M2 and to only one coronal plane, both ipsi- and contralaterally. Labelling patterns following PVO injection was similar to those observed following ALO injection, although resulting in weaker labelling in contralateral M1. Overall, input from motor cortex suggests both an anterior-posterior difference, and a medial-lateral difference such that PLO receives much weaker motor inputs than ALO and PVO (Table 2).

#### Sensory cortex

Following CTB injection into PVO, but not ALO or PLO, we observed weak labelling present in the lateral, mediolateral, and mediomedial areas of the secondary visual cortex, and weak to moderate labelling in the barrel cortex and in the forelimb, hindlimb, and trunk regions of the primary somatosensory cortex. In contrast, except for weak labelling in the barrel cortex after ALO injection, we observed no labelled cells in the visual or the somatosensory cortices following injection into lateral OFC (either ALO or PLO). Following injection into either ALO, PLO, or PVO, we consistently detected moderate labelling in piriform cortex and weak labelling in granular insular cortex. However, posterior to −1.88 mm from bregma, the piriform cortex exhibited only weak labelling from PVO injections. The auditory cortex was devoid of labelled cells following injection into either subdivision. Therefore, there was strong evidence of primary multi-sensory inputs into PVO, but not into ALO or PLO (Table 2).

### Thalamus

Detailed schematics for the thalamus are shown in Figures 4 and 5. ALO, PLO, and PVO, each exhibit distinct connectivity patterns with the mediodorsal (MD) and submedius (Sub) nuclei of the thalamus.

**Figure 4.**
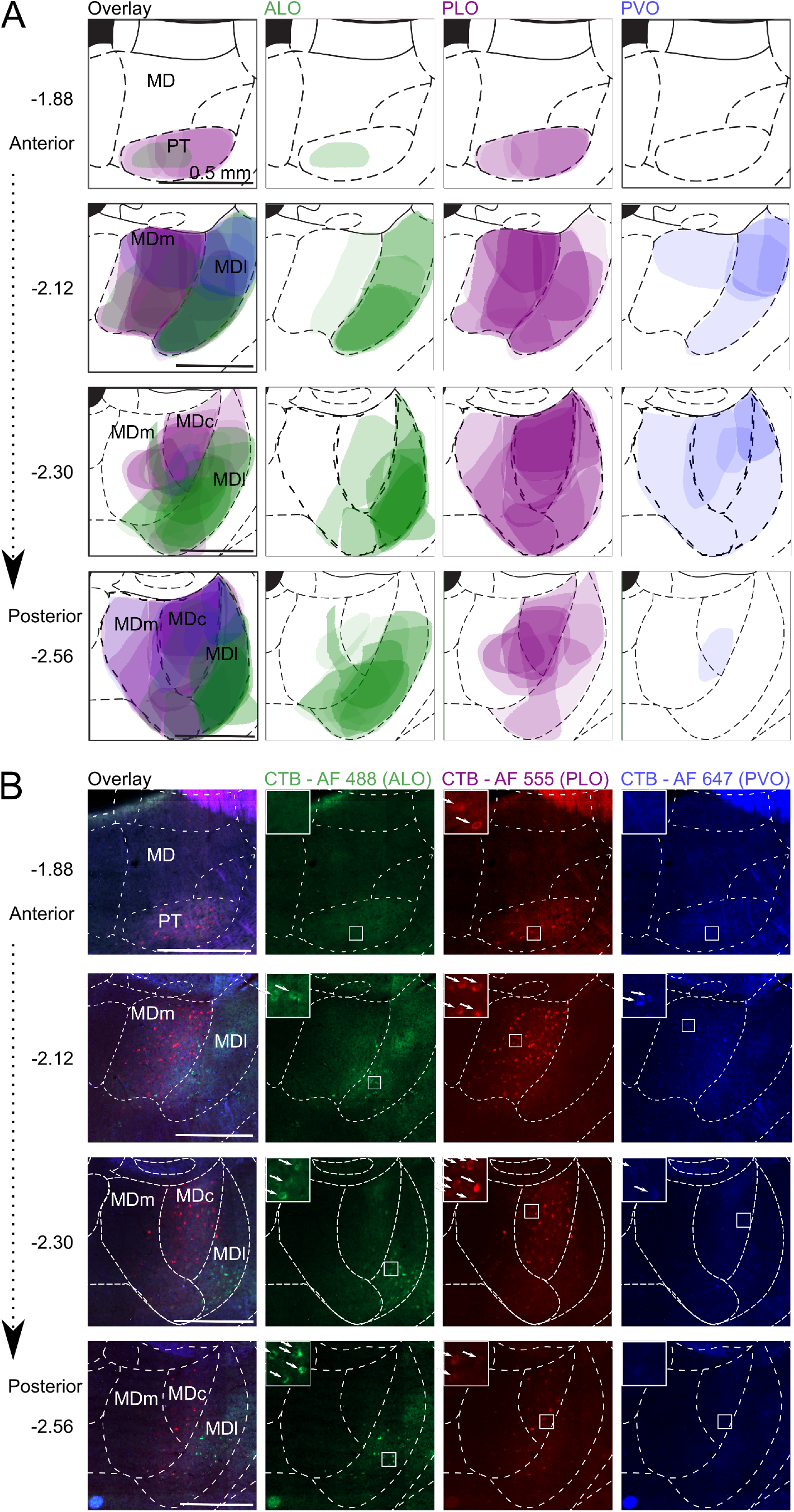
Localisation and density of retrogradely-labelled cells in the MD following CTB injection into ALO, PLO, or PVO. (A) Shaded areas correspond to superimposed hand drawings of the labelling in each brain included in the analysis, using an opacity value proportional to the labelling density level. *n* ALO = 6, *n* PLO = 5, *n* PVO = 4 injections. (B) Micrographs from representative brain with successful triple labelling. c: central; l: lateral; m: medial; AF: Alexa Fluor; ALO: lateral orbitofrontal cortex, anterior area; CTB: cholera toxin subunit B; MD: mediodorsal nucleus of the thalamus; PLO: lateral orbitofrontal cortex, posterior area; PVO: ventral orbitofrontal cortex, posterior area.

**Figure 5.**
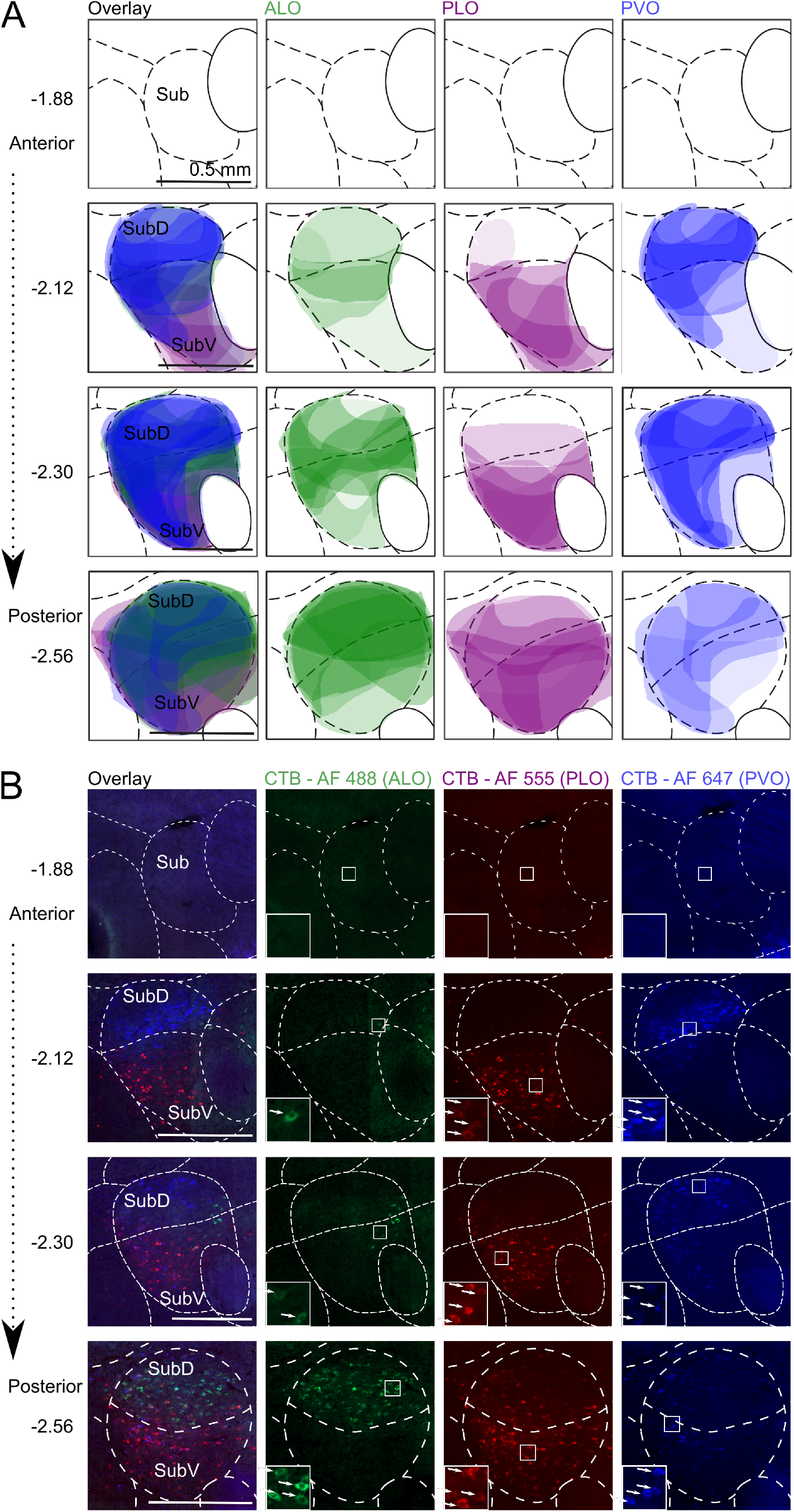
Localisation and density of retrogradely-labelled cells in Sub following CTB injection into ALO, PLO, or PVO. (A) Shaded areas correspond to superimposed hand drawings of the labelling in each brain included in the analysis, using an opacity value proportional to the labelling density level. *n* ALO = 6, *n* PLO = 5, *n* PVO = 4 injections. (B) Micrographs from representative brain with successful triple labelling. D: dorsal; V: ventral; AF: Alexa Fluor; ALO: lateral orbitofrontal cortex, anterior area; CTB: cholera toxin subunit B; PLO: lateral orbitofrontal cortex, posterior area; PVO: ventral orbitofrontal cortex, posterior area; Sub: submedius nucleus of the thalamus.

#### Mediodorsal nucleus

ALO receives strongest projections from the lateral part (MDl), particularly its most ventral portion. PLO receives strong projections from both the medial (MDm) and central (MDc) parts of the medial dorsal thalamic nucleus, and only weak ones from MDl (Figure 4). In contrast, PVO receives only light projections from either lateral, medial or central MD.

Supporting the observed pattern of labelling statistically (summarised in Figure 6-A), OFC subregions received distinct patterns of projections from MD (significant main effect of MD thalamus portion, *F*_(2, 30.2)_ = 4.32, *p* = .022; and injection site x MD thalamus portion interaction, *F*_(4, 30.3)_ = 3.80, *p* = .013). ALO receives stronger projections from MDl than from MDm (ALO: MDl vs MDm, *t*_(36)_ = 3.40, *p* = .005) or MDc (ALO: MDl vs MDc, *t*_(36)_ = 2.95, *p* = .017), but similar projection strength from MDc and MDm (ALO: MDc vs MDm, *t*_(36)_ = 0.45, *p* = .956). In contrast to ALO, there were no differences in the relative strength of projections from MD subregions to PLO and PVO (all *t_(36)_* < |1.82|, *p* > .078). There was a main effect of injection site (ALO, PLO, PVO; *F*_(2, 34.1)_ = 16.03, *p* < .001), with PLO receiving relatively strong projections and PVO receiving relatively weak projections from MD overall (PLO vs PVO: *t*_(32.6)_ = 5.60, *p* < .001; PLO vs ALO: *t*_(35.6)_ = 3.47, *p* = .004; PVO vs ALO: *t*_(34.0)_ = −2.27, *p* = .086).

**Figure 6.**
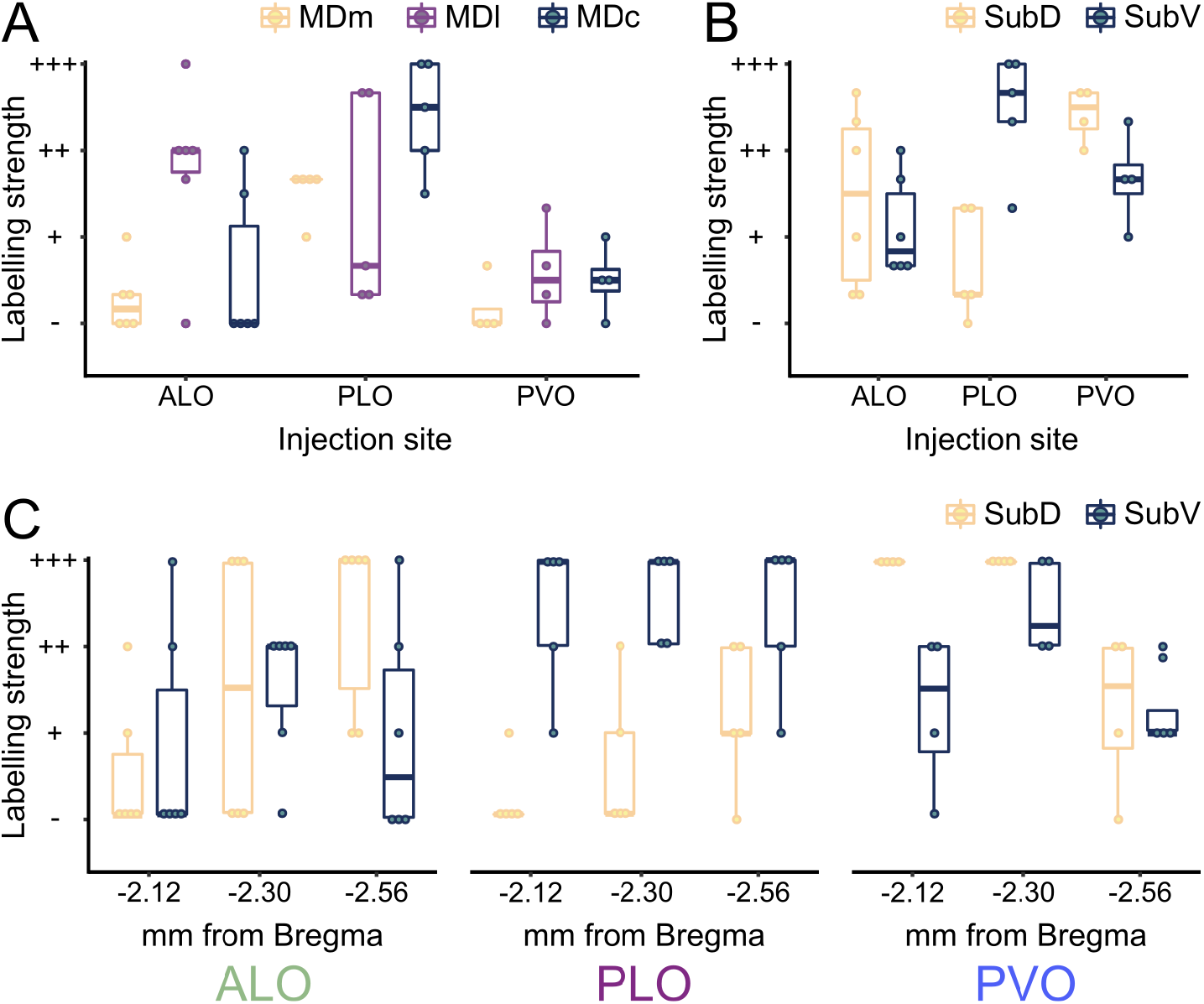
Labelling strength in MD and Sub following injection of CTB into ALO, PLO, or PVO. (A) Average labelling strength in MD across coronal sections −1.88, −2.12, −2.30, and −2.56 mm from bregma.) (B) Average labelling strength in Sub across coronal sections −2.12, −2.30, and −2.56 mm from bregma. (C) Labelling strength in Sub after ALO, PLO, or PVO injections at coronal sections −2.12, −2.30, or −2.56 mm from bregma. *n* ALO = 6, *n* PLO = 5, *n* PVO = 4 injections. c : central; l: lateral; m: medial; D: dorsal; V: ventral; ALO: lateral orbitofrontal cortex, anterior area; CTB: cholera toxin subunit B; MD: mediodorsal nucleus of the thalamus; PLO: lateral orbitofrontal cortex, posterior area; PVO: ventral orbitofrontal cortex, posterior area; Sub: submedius nucleus of the thalamus.

#### Submedius nucleus

All portions of Sub project to every region of the OFC investigated here, although there is a clear spatial segregation in relation to where in Sub the strongest projections originate (Figure 5). ALO receives strongest projections from the posterior part of the dorsal submedius nucleus (SubD). In contrast, most afferents from Sub to PLO come from the ventral portion (SubV). On the other hand, PVO receives most Sub projections from the most anterior half, prominently from the dorsal portion, and slightly less from the ventral part.

First, we quantified the overall pattern of projections from Sub (Figure 6-B), focusing on the average labelling strength across coronal sections (averaging across −2.12, −2.38, and −2.56 mm from bregma). This revealed that PLO receives stronger projections from SubV than SubD (PLO: SubV vs SubD: *t*_(24)_ = 4.13, *p* = .001), whereas ALO and PVO (ALO: SubV vs SubD: *t*_(24)_ = −0.82, *p* = .804; PVO: SubV vs SubD: *t*_(24)_ = −1.59, *p* = .330) do not appear to be differentiated (supported by an injection site x Sub portion interaction, *F*_(2, 18.2)_ = 13.36, *p* < .001). Furthermore, PVO receives stronger overall projections from Sub than ALO (main effect of injection site *F*_(2, 20.2)_ = 7.09, *p* = .005; PVO vs ALO, *t*_(20)_ = 3.77, *p* = .004).

Next, we explored the differences in the topographical organisation of these projections along the anterior-posterior axis (Figure 6-C). A significant 3-way interaction of injection site x Sub portion x coronal section (*F*_(4, 66.1)_ = 3.27, *p* = .016), indicated that the distribution and strength of projections from Sub along the anterior-posterior axis differed depending on the orbitofrontal injection site. We explored this 3-way interaction by performing separate coronal section (−2.12, −2.30, −2.56 mm from bregma) x Sub portion (SubV, SubD) ANOVAs for each injection site (ALO, PLO, PVO).

Projections to ALO increased from anterior to posterior Sub (main effect of coronal section, *F*_(2, 25)_ = 4.08, *p* = 0.029; coronal sections −2.12 vs −2.30 mm: *t*_(25)_ = −1.98, *p* = .166; −2.12 vs −2.56 mm: *t*_(25)_ = −2.77, *p* = .031; −2.30 vs −2.56 mm: *t*_(25)_ = −0.79, *p* = .820), but were not differentiated by dorsal or ventral regions within Sub (main effect of Sub portion: SubV vs SubD, *F*_(1, 25)_ = 0.34, *p* = .564; coronal section x Sub portion interaction, *F*_(2, 25)_ = 1.88, *p* = .174). In contrast, Sub projections to PLO were not differentiated along the anterior-posterior axis (main effect of coronal section: *F*_(2, 20)_ = 1.84, *p* = .185; coronal section x Sub portion interaction: *F*_(2, 20)_ = 2.00, *p* = .161), but labelling was significantly stronger in SubV than in SubD (main effect of Sub portion, SubV vs SubD: *F*_(1, 20)_ = 74.09, *p* < .001).

Finally, the relative strength of projections to PVO originating from SubD and SubV changed along the anterior-posterior axis (coronal section x Sub portion interaction (*F*_(2, 15)_ = 5.82, *p* = .013). Stronger labelling was observed in SubD than in SubV at −2.12 mm from bregma (−2.12 mm: SubD vs SubV, *t*_(18)_ = 4.88, *p* < .001), but no differences were found between labelling in the Sub portions at more posterior coronal sections (−2.30 mm: SubD vs SubV, *t*_(18)_ = 1.68, *p* = .298; −2.56 mm: SubD vs SubV: *t*_(18)_ = 2.46, *p* = .993). Overall projections from Sub to PVO were strongest at −2.30 mm from bregma (main effect of coronal section, *F*_(2, 15)_ = 17.76 *p* < .001; −2.12 vs −2.30 mm, t_(15)_ = −2.81, *p* = .039; −2.30 vs −2.56 mm, t_(15)_ = 5.96, *p* < .001; −2.12 vs −2.56 mm, t_(15)_ = 3.15, *p* = .020). Overall, this pattern of projections suggests an anterior-posterior gradient of projections from Sub to ALO, projections from Sub to PLO originate predominantly in SubD, and a topographically unique pattern of projections to PVO that differs across anterior-posterior and dorsal-ventral axes.

#### Paratenial nucleus

The paratenial thalamic nucleus (PT; Figure 4) consistently exhibited moderate labelling following PLO injection but very light to no labelling with injections into ALO or PVO.

#### Nucleus reuniens

In the reuniens thalamic (Re) nucleus, we observed light labelling following tracer injections into both lateral and ventral posterior OFC, but mostly absent after injections into ALO (Figure 2).

In sum, the thalamus sends distinct and topographically organised projections from MD and Sub into ALO, PLO and PVO (Figure 6, Table 2). Here, the anterior-posterior distinction between ALO and PLO is as clear and differentiated as the medial-lateral distinction between PLO and PVO. Projections from Re to PLO and PVO, but not to ALO, reinforce the anterior-posterior distinction within LO. Labelling in PT after PLO injection but not after ALO or PVO also reinforces this anterior-posterior distinction within LO as well as a medial-lateral distinction between PLO and PVO.

### Temporal lobe

Lateral entorhinal cortex exhibited weak labelling following injection into either ALO or PLO but stronger labelling following PVO injection. Labelling was also detected in the perirhinal cortex after tracer injection into PVO but was not consistently present following ALO or PLO injections (see Figure S3 and S4). In summary, although labelling in the entorhinal and perirhinal was not always consistent, the labelling patterns observed suggest a gradient between their inputs into LO and VO.

#### Amygdala

A detailed schematic for the amygdala is shown in Figure 7 (see Figure S7 for representative histological images). The anterior part of the basolateral amygdala (BLA) exhibited moderate labelling following PLO injection but only weak labelling after either ALO or PVO injections (Figure 7, Figure S7). We observed a similar pattern in the dorsolateral part of the lateral amygdala (LaDL), with the difference that it did not contain any labelled cells following PVO injection. We only detected weak labelling in the posterior part of the basolateral amygdala (BLP) and in the ventromedial (LaVM) and ventrolateral (LaVL) parts of the lateral amygdala following tracer injection into PLO. There were no labelled cells outside of the basolateral and lateral regions of the amygdala. Overall, amygdala labelled cells were most prevalent in BLA and following injection into PLO (Table 2), suggesting both an anterior-posterior gradient within LO and a medial-lateral gradient between PLO and PVO.

**Figure 7.**
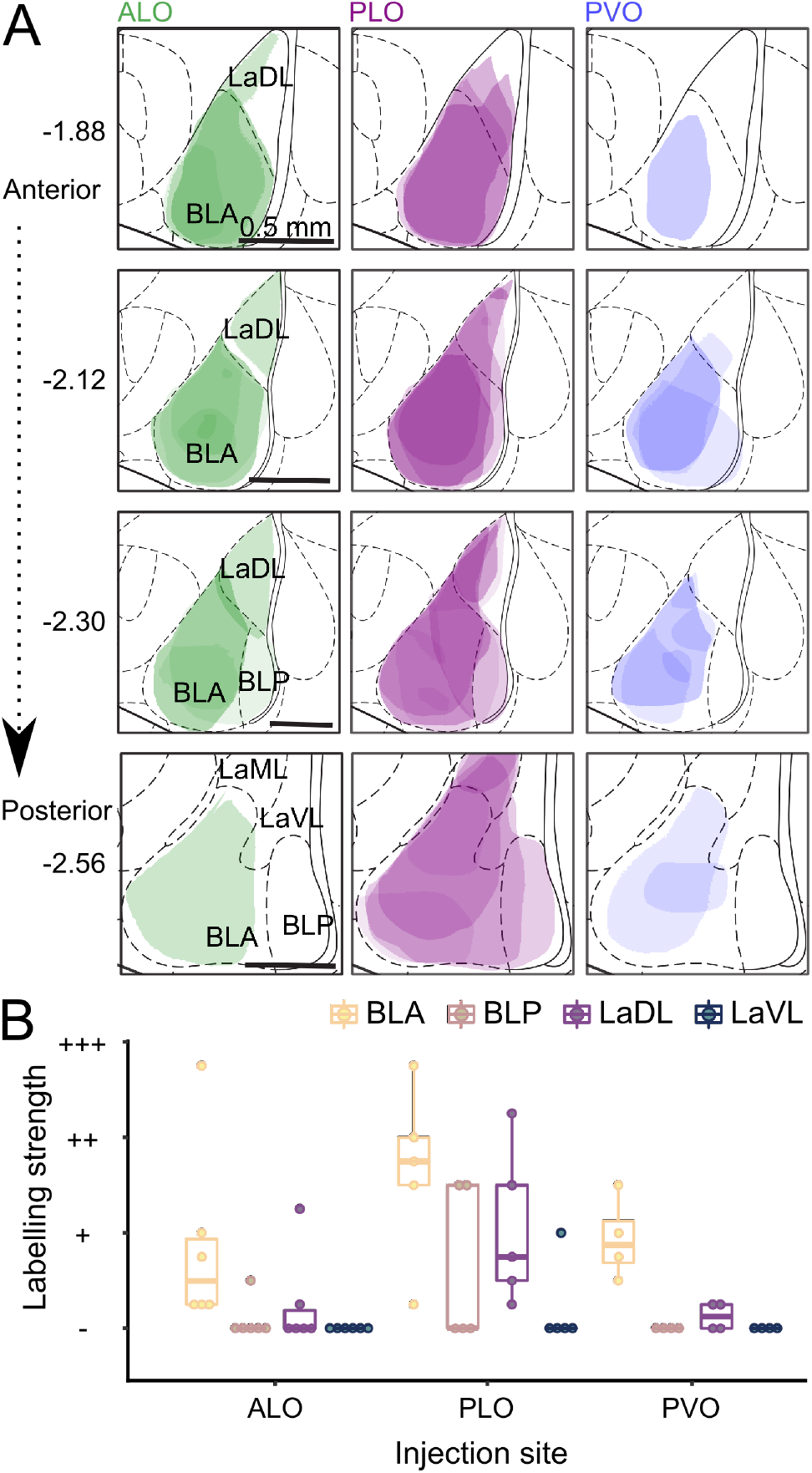
Labelling in the amygdala following CTB injection into ALO, PLO, or PVO. (A) Shaded areas correspond to superimposed hand drawings of the labelling in each brain included in the analysis, using an opacity value proportional to the labelling density level. *n* ALO = 6, *n* PLO = 5, *n* PVO = 4 injections. (B) Average labelling strength in regions of the amygdala across coronal sections −1.88, −2.12, −2.30 and −2.56 mm from bregma. *n* ALO = 6, *n* PLO = 5, *n* PVO = 4 injections. ALO: lateral orbitofrontal cortex, anterior area; CTB: cholera toxin subunit B; BLA: basolateral amygdala, anterior part; BLP: basolateral amygdala, posterior part; LaDL: lateral amygdala, dorsolateral part; LaML: lateral amygdala, mediolateral part; LaVL, lateral amygdala, ventrolateral part; PLO: lateral orbitofrontal cortex, posterior area; PVO: ventral orbitofrontal cortex, posterior area.

Supporting these observations, statistical analysis of the average labelling strength across coronal sections (Figure 7-B) revealed a main effect of injection site (ALO, PLO, PVO; *F*_(3, 47.1)_ = 8.38, *p* < .001), with PLO receiving stronger amygdala projections than ALO (PLO vs ALO: *t*_(47.9)_ = 3.73, *p* = .002) or PVO (PLO vs PVO: *t*_(45.8)_ = 3.30, *p* = .006), and ALO and PVO receiving similar strength of projections (ALO vs PVO: *t*_(47.4)_ = −0.23, *p* = .994). There was also a main effect of amygdala portion (BLA, BLP, LaVL, LaVM; *F*_(3, 42.4)_ = 15.76, *p* < .001), with BLA sending the strongest projections to OFC (BLA vs BLP: *t*_(42.4)_ = 4.40, *p* < .001; BLA vs LaVL: *t*_(42.4)_ = 4.40, *p* < .001; BLA vs LaDL: *t*_(42.4)_ = 2.68, *p* = .061), and LaDL sending stronger projections than LaVL (LaDL vs LaDL: *t*_(42.4)_ = 3.97, *p* = .002). However, the strength of projections from these amygdala portions did not differentially project to OFC subregions (amygdala portion x injection site interaction, *F*_(6, 42.4)_ = 1.84, *p* = .115).

## Discussion

Here, we tested the hypothesis that OFC functional heterogeneity predicts meaningful differences in connectivity by simultaneously characterising the patterns of inputs of the anterior and posterior portions of LO, which have been previously found to be functional distinct (Panayi & Killcross, 2018). Additionally, we contrast the input patterns into ALO and PLO with inputs into PVO, an anatomically adjacent OFC portion which is thought to be functionally distinct (Balleine et al., 2011; Corwin et al., 1994). Our approach allowed us to assess the extent of overlap, convergence and divergence between the projections of these OFC subdivisions, and thus define their topographic relationships. By using the same tracer to compare the connectivity across the subdivisions, we minimised potential interpretation problems caused by variability in uptake or transport by the cells, spread in the tissue or extent of local necrosis.

The OFC receives its main inputs from the amygdala, thalamic nuclei, periaqueductal gray, and midbrain dopamine neurons (e.g. Krettek & Price, 1977b; Mcdonald, 1998; Murphy & Deutch, 2018; Ongur & Price, 2000). Our neuroanatomical characterisation revealed substantial differences in cortical and sub-cortical inputs into ALO, PLO, and PVO. Specifically, we identified robust differences in the topographic organisation and in the gradation of connectivity strength into the OFC subdivisions investigated. Such differences were observed in the inputs from the medial prefrontal cortex, motor cortex, sensory cortices, amygdala, and thalamus (Figure 8). We did not assess midbrain inputs to OFC in our study. However, there are moderate dopaminergic projections from midbrain, including ventral tegmental area, dorsal raphe nucleus and ventral periaqueductal gray to posterior LO and laterally adjacent AI (Murphy & Deutch, 2018), as well as noradrenergic projections from the locus coeruleus to LO, VO and MO (Cerpa et al., 2019) and to AI (Gerfen & Clavier, 1979).

**Figure 8.**
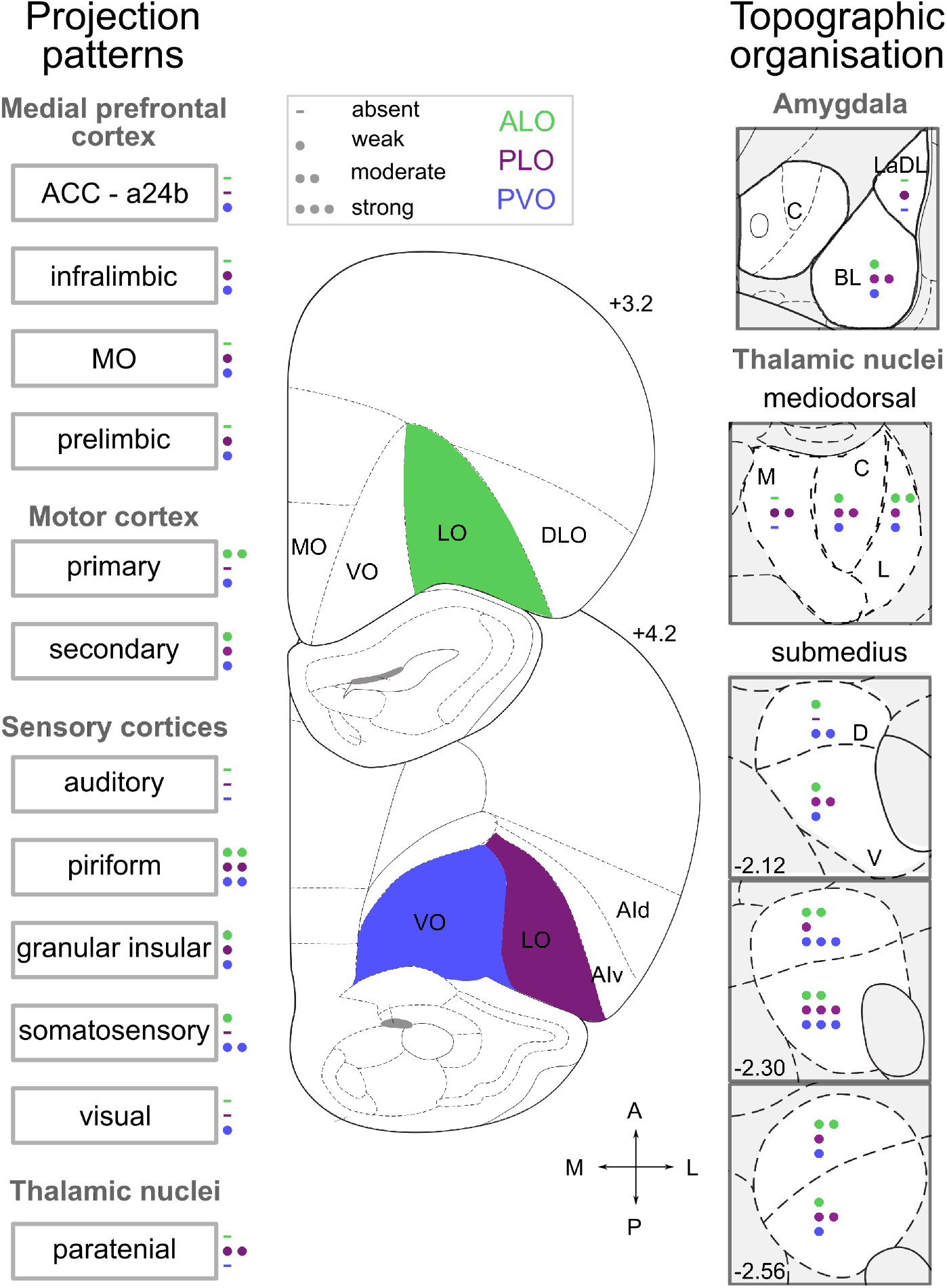
Density of key inputs into ALO, PLO and PVO. The different number of circles represent the average density of retrogradely-labelled cells following CTB injection (i.e. absent, weak, moderate or strong) into ALO, PLO, and PVO. Panels depicting labelling in amygdala and in MD thalamus represent the average density labelling in all sections quantified and not at that particular coronal section. Distances shown are distances from bregma in mm. a24b: anterior cingulate cortex, area 24b; ACC: anterior cingulate cortex; AId: agranular insular cortex, dorsal part; AIv: agranular insular cortex, ventral part; ALO: lateral orbitofrontal cortex, anterior part; CTB: cholera toxin subunit B; DLO: orbitofrontal cortex, dorsolateral part; IL: infralimbic cortex; LaDL: lateral amygdala, dorsolateral part; LO: orbitofrontal cortex, lateral part; M1: primary motor cortex; M2: secondary motor cortex; MO: orbitofrontal cortex, medial part; PL: prelimbic cortex; PLO: lateral orbitofrontal cortex, posterior part; PVO: ventral orbitofrontal cortex, posterior part; VO: orbitofrontal cortex, ventral part; A: anterior; P: posterior; BL: basolateral; C: central; M: medial; L: lateral; D: dorsal; V: ventral.

### Distinct and topographically organised thalamic inputs to OFC subdivisions

Thalamo-cortical connectivity has historically been one of the key criteria used to both segregate cortical regions and define functional circuits (Alexander, 1986; Rose & Woolsey, 1948). Medial, lateral, and central nuclei of the MD thalamus receive partially overlapping but distinct cortical and subcortical afferents (Groenewegen, 1988). In our study, retrogradely-labelled cells in MD thalamus showed a notably separate pattern of projections into ALO, PLO, and PVO. In fact, the distinction between ALO and PLO was just as clear as the medial-lateral distinction between PLO and PVO.

The Sub shares strong reciprocal connections with the OFC, especially with VO/LO (Alcaraz et al., 2015; Coffield et al., 1992; Kuramoto et al., 2017; Reep et al., 1996; Yoshida et al., 1992). In our tracing study we observed the most pronounced labelling in this area, with all portions of this thalamic nucleus projecting to every region of the OFC that we investigated (Figures 4, 6). Again, rather than a uniform connectivity pattern, we observed clear spatial segregation in relation to where in the Sub the strongest projections originate. The distinct medial-lateral gradient of OFC inputs from Sub we observed has been previously reported (Alcaraz et al., 2015; Reep et al., 1996). However, we also identified a novel pattern of anterior-posterior projections from Sub which provides the clearest evidence of a distinction in afferents across the three OFC subregions we investigated in our study: ALO, PLO, and PVO. Our results revealed that PVO receives most Sub projections from the most anterior half of the Sub, especially from the dorsal portion, and slightly less from the ventral part. Intriguingly, results from previous tracing studies (Kuramoto et al., 2017; Reep et al., 1996) revealed that AVO receives inputs from SubV, suggesting a potential anterior-posterior distinction might also be present here. Such distinction within VO is also supported by stronger noradrenergic projections from the locus coeruleus to PVO than to AVO (Cerpa et al., 2019).

The OFC is the main recipient of Sub projections and, in addition to the subdivisions already mentioned, Sub also sends projections to MO (Kuramoto et al., 2017; Yoshida et al., 1992). Most of the double- or triple-labelled cells we observed were present in the Sub. This means that, unlike other regions studied here, a proportion of information from Sub neurons is broadcast to ALO, PLO, and PVO. It has previously been reported there are neurons in this region that simultaneously project to lateral and ventral OFC, potentially linking their functions under certain conditions (Kuramoto et al., 2017). Other than OFC inputs, the Sub receives mainly sensory and pain modulation information (Yoshida et al., 1992). The strong VO/LO-Sub reciprocal connection seems to be involved in a pain modulation pathway (Coffield et al., 1992; Kuramoto et al., 2017; Tang et al., 2009). While little is known about Sub function, recent studies exploring the functional significance of its connectivity with the OFC, revealed that an intact Sub is necessary for updating both stimulus-outcome (Alcaraz et al., 2015) and action-outcome associations (Fresno et al., 2019).

We also observed distinct labelling patterns in the paratenial thalamic nucleus, which consistently exhibited moderate labelling following PLO injection but very light to no labelling with injections into ALO (Figure 2; Table 2). Following injections of anterograde tracer into PT, Vertes & Hoover (2008) observed a similar distinction in its projections to the anterior and posterior portions of LO. Functional studies investigating PT-OFC interaction might help understand the significance of this OFC anterior-posterior distinction.

### Multisensory nature of OFC inputs

The lateral orbital network, comprised of VO, LO, and AI, is sometimes also referred to as a sensory network because of its strong multi-modal sensory inputs (Ongur & Price, 2000; Price, 2007). Here, we observed evidence of multi-sensory inputs into PVO. This in agreement with the attentional account of VO proposed by Hoover & Vertes (2011) and the connectivity described by Corwin & Reep (1998), showing that VO receives direct visual and somatosensory but not auditory information. By contrast, primary sensory inputs into both ALO and PLO, other than gustatory and olfactory, are sparse or absent. While the OFC is often considered a site of multi-sensory input, these results highlight a medial-lateral distinction in the diversity of primary sensory inputs into LO and VO.

### Amygdala input patterns into OFC subdivisions

OFC and BLA share strong reciprocal connections and their projections are topographically organised in both directions (Mcdonald, 1991; Mcdonald, 1998; Reep et al., 1996). In our tracing experiments, amygdala labelled cells were most prevalent following injection into PLO, originating most strongly in anterior BLA. By contrast, we observed only weak labelling after either ALO or PVO injections. The posterior OFC medial-lateral difference is consistent with the results obtained by Kita & Kitai (1990) following injections of anterograde tracer into anterior BLA: dense labelling in PLO but not in PVO. OFC function has been closely linked to that of the amygdala, in particular its basolateral nucleus. Both OFC and BLA have been implicated in processes that support outcome-guided behaviour through associative learning of sensory-specific representations and predictive-cues, with lesions of either region impairing performance on outcome devaluation and reversal learning tasks (for reviews see Balleine & Killcross, 2006; Sharpe & Schoenbaum, 2016). The amygdala has been implicated specifically in tracking previous outcomes and comparing them to current outcomes (Izquierdo et al., 2013; Lichtenberg et al., 2017; Schoenbaum et al., 2003; Stalnaker et al., 2007). Interestingly, the stronger inputs from BLA to PLO we observed shed light on previously observed functional differences between ALO and PLO: posterior but not anterior LO lesions disrupt reversal learning, suggesting PLO is critical for updating the value of expected outcomes (Panayi & Killcross, 2018). Moreover, close inspection of a recent paper showing the necessity of OFC inputs from BLA for using cue-generated reward expectations to guide decision making show that these mainly terminated in PLO (Lichtenberg et al., 2017).

Paralleling the observations by Panayi & Killcross (2018), a study in non-human primates found that posterior OFC (area 13) is required for updating outcome valuations during a selective satiety devaluation procedure, whereas anterior OFC (area 11) is necessary for translating this knowledge into action during goal selection (Murray et al., 2015). Stronger inputs from BLA to posterior (area 13a) than to anterior OFC (area 11) have been previously observed in primates (Amaral & Price, 1984). This reinforces the idea, which had been previously hinted at by these functional studies (Murray et al., 2015; Panayi & Killcross, 2018), of similar functional organisation principles between rodent and primate OFC subdivisions when functional differences along the anterior-posterior axis are considered.

### Anterior and posterior LO are part of distinct anatomical and functional circuits

While both PVO and PLO receive projections from regions along the medial wall, including IL, PL, and MO (Figure 2; Table 2), such projections are mostly absent following injection into ALO. This is consistent with previous reports studying inputs from PL and IL to the OFC, which also suggest anterior-posterior differences (Takagishi & Chiba, 1991; Vertes, 2004). Even though PVO and PLO injections were in the same anteroposterior plane and PVO is situated more medially, there were comparable degrees of labelling along the medial wall following injections into either of the two subdivisions. This argues against issues with polysynaptic labelling or tracer spreading within the OFC, especially between PVO and PLO injections. Indeed, we observed that CTB deposits that extended into adjacent AI produced a clearly distinct pattern of inputs to the adjacent PLO, with dense innervation from PL, MO and BLA (see Figure S6; consistent with Murphy & Deutch, 2018; Reep et al., 1996).

A number of studies have proposed a functional dissociation between OFC and PL in supporting Pavlovian and instrumental conditioning, respectively (Balleine & Dickinson, 1998; Corbit & Balleine, 2003; Ostlund & Balleine, 2005; Ostlund & Balleine, 2007). Both PL and OFC share strong reciprocal connections with the BLA (Mcdonald, 1991; Mcdonald, 1998), which in turn has been shown to play a role in both Pavlovian and instrumental outcome-mediated decision making (Balleine & Killcross, 2006; Johnson et al., 2009). Therefore, the similar distinction in the density of projections from PL and BLA to anterior and posterior LO, suggest that BLA®PLO and PL®PLO might form a functional circuit involved in assessing the motivational significance of actions and stimuli, respectively.

#### Functional implications

Even very functionally distinct brain regions often show substantial overlap in their inputs and outputs (Passingham et al., 2002). It is, thus, not surprising that different portions of the OFC show the graded connectivity patterns we described here. It might, therefore, be useful to consider the question of what makes two adjacent regions distinct. We believe that integrating neuroanatomical characterisation with functional differences is essential. For instance, VO and LO are generally considered separate regions (Krettek & Price, 1977; Price, 2006; Ray & Price, 1992). This distinction is supported by functional distinctions (e.g. Balleine et al., 2011; Izquierdo, 2017) and differences in efferent (e.g. Schilman et al., 2008) and afferent (Table 2) projections.

In light of this standard distinction between VO and LO, we ought to consider a division between the anterior and posterior portions of LO. A functional dissociation between ALO and PLO has been previously reported (Panayi & Killcross, 2018) and our neuroanatomical characterisation of their inputs has shed light on the circuitry underlying those differences. We observed a similar degree of distinction between the afferent projections into ALO and PLO and the projections into PLO and PVO. Thus, while further studies might be necessary, the evidence so far firmly supports a distinction along the anterior-posterior axis of LO. We did not explore whether a similar division might exist within VO. However, the projection patterns we observed in the submedius nucleus of the thalamus following injection into PVO differ substantially from those seen by Kuramoto and colleagues (2017) following injection of the same retrograde tracer into AVO.

Our results revealed that different portions of the Sub send distinct, strong and topographically organised projections to OFC subdivisions. Thus, the Sub presents itself as a key anatomical area within the thalamus whose projections may allow the distinction of orbital subregions more effectively than those of the MD thalamus, which have traditionally been used when defining prefrontal circuits (Rose & Woolsey, 1948). The specific pattern and relative density of anatomical projections between brain regions is also one of the criteria used to establish homology between cortical areas of different species (Rose & Woolsey, 1948; Uylings et al., 2003). A detailed understanding of the anatomy of OFC subdivisions will, therefore, allow for connectivity-based inferences on cross species-homologies. Thus, further anatomical and functional characterisation of OFC subdivisions will likely be key to establish clearer homologies between rodent and primate OFC.

## Authors contributions

MCP and MEW conceived the project. IVB and MCP conducted the experiments. IVB analysed the data, with input from MCP and MEW. IVB, MCP and MEW wrote the paper.

## Declarations of interest

None.

## Acknowledgments

This work was supported by a Biotechnology and Biological Sciences Research Council Doctoral Training Partnership studentship awarded to IVB; and by a Wellcome Senior Research Fellowship awarded to MEW (202831/Z/16/Z). The authors thank Allan Wainman from the Micron Advanced Bioimaging Unit (supported by Wellcome Strategic Awards 091911/B/10/Z and 107457/Z/15/Z) for support and assistance in this work.

## Abbreviations

a24b: anterior cingulate cortex, area 24b
AI, d, v, P: agranular insular cortex, dorsal, ventral, posterior parts
ALO: lateral orbitofrontal cortex, anterior part
Au1: auditory cortex, primary area
AVO: ventral orbitofrontal cortex, anterior part
BLA: basolateral nucleus of amygdala
BLP: basolateral nucleus of the amygdala, posterior part
DLO: dorsolateral orbitofrontal cortex
GI: granular insular cortex
IL: infralimbic cortex
LaDL: lateral amygdala, dorsolateral part
LaVL: lateral amygdala, ventrolateral part
LaVM: lateral amygdala, ventromedial part
LEnt: lateral entorhinal cortex
LO: lateral orbitofrontal cortex
M1: primary motor cortex
M2: secondary motor cortex
MD, m, c, l: mediodorsal nucleus of the thalamus, medial, central, lateral parts
MO: medial orbitofrontal cortex
Pir: piriform cortex
PL: prelimbic cortex
PLO: lateral orbitofrontal cortex, posterior part
PRh: perirhinal cortex
PT: paratenial nucleus of the thalamus
PVO: ventral orbitofrontal cortex, posterior part
Re: nucleus reuniens of the thalamus
RS: Retrosplenial cortex
S1BF: primary somatosensory cortex, barrel field
S1J: primary somatosensory cortex, jaw region
S1HL/FL: primary somatosensory cortex, hindlimb and forelimb regions
S1Tr: primary somatosensory cortex, trunk region
S2: secondary somatosensory cortex
Sub, D, V: submedius nucleus of the thalamus, dorsal, ventral parts
V2: secondary visual cortex
VO: ventral orbitofrontal cortex

## Notes

### Competing Interest Statement

The authors have declared no competing interest.

### Summary of Updates

In this version we: -Reorganised some of the figures and improved their quality/resolution. -Added a new figure obtained with a confocal microscope depicting example single-, double-, and triple-labelled cells in the submedius nucleus of the thalamus (Figure 3). -Performed statistical analyses on the labelling levels observed in the amygdala, and the mediodorsal and submedius nuclei of the thalamus.

## References

Alcaraz, F., Marchand, A. R., Vidal, E., Guillou, A., Faugère, A., Coutureau, E., & Wolff, M. (2015). Flexible use of predictive cues beyond the orbitofrontal cortex: Role of the submedius thalamic nucleus. Journal of Neuroscience, 35(38), 13183–13193. https://doi.org/10.1523/JNEUROSCI.1237-15.2015

Alexander, G. (1986). Parallel Organization of Functionally Segregated Circuits Linking Basal Ganglia and Cortex. Annual Review of Neuroscience, 9(1), 357–381. https://doi.org/10.1146/annurev.neuro.9.1.357

Amaral, D. G., & Price, J. L. (1984). Amygdalo-cortical projections in the monkey (Macaca fascicularis). The Journal of Comparative Neurology, 230(4), 465–496. https://doi.org/10.1002/cne.902300402

Balleine, B. W., & Dickinson, A. (1998). Goal-directed instrumental action: Contingency and incentive learning and their cortical substrates. Neuropharmacology, 37(4–5), 407–419. https://doi.org/10.1016/S0028-3908(98)00033-1

Balleine, B. W., & Killcross, S. (2006). Parallel incentive processing: an integrated view of amygdala function. Trends in Neurosciences, 29(5), 272–279. https://doi.org/10.1016/j.tins.2006.03.002

Balleine, B. W., Leung, B. K., & Ostlund, S. B. (2011). The orbitofrontal cortex, predicted value, and choice. Annals of the New York Academy of Sciences, 1239(1), 43–50. https://doi.org/10.1111/j.1749-6632.2011.06270.x

Barreiros, I. V., Ishii, H., Walton, M. E., & Panayi, M. C. (2020). Defining an orbitofrontal compass: functional and anatomical heterogeneity across anterior-posterior and medial-lateral axes. Submitted.

Bradfield, L. A., & Hart, G. (2020). Rodent medial and lateral orbitofrontal cortices represent unique components of cognitive maps of task space. Neuroscience and Biobehavioral Reviews, 108(November 2019), 287–294. https://doi.org/10.1016/j.neubiorev.2019.11.009

Bradfield, L. A., Hart, G., & Balleine, B. W. (2018). Inferring action-dependent outcome representations depends on anterior but not posterior medial orbitofrontal cortex. Neurobiology of Learning and Memory, 155(August), 463–473. https://doi.org/10.1016/j.nlm.2018.09.008

Cerpa, J. C., Marchand, A. R., & Coutureau, E. (2019). Distinct regional patterns in noradrenergic innervation of the rat prefrontal cortex. Journal of Chemical Neuroanatomy, 96(November 2018), 102–109. https://doi.org/10.1016/j.jchemneu.2019.01.002

Coffield, J. A., Bowen, K. K., & Miletic, V. (1992). Retrograde tracing of projections between the nucleus submedius, the ventrolateral orbital cortex, and the midbrain in the rat. Journal of Comparative Neurology, 321(3), 488–499. https://doi.org/10.1002/cne.903210314

Conte, W. L., Kamishina, H., & Reep, R. L. (2009). Multiple neuroanatomical tract-tracing using fluorescent Alexa Fluor conjugates of cholera toxin subunit B in rats. Nature Protocols, 4(8), 1157–1166. https://doi.org/10.1038/nprot.2009.93

Corbit, L. H., & Balleine, B. W. (2003). The role of prelimbic cortex in instrumental conditioning. Behavioural Brain Research, 146(1–2), 145–157. https://doi.org/10.1016/j.bbr.2003.09.023

Corwin, J. V., Fussinger, M., Meyer, R. C., King, V. R., & Reep, R. L. (1994). Bilateral destruction of the ventrolateral orbital cortex produces allocentric but not egocentric spatial deficits in rats. Behavioural Brain Research, 61(1), 79–86. https://doi.org/10.1016/0166-4328(94)90010-8

Corwin, J. V., & Reep, R. L. (1998). Rodent posterior parietal cortex as a component of a cortical network mediating directed spatial attention. Psychobiology, 26(2), 87–102. https://doi.org/10.3758/BF03330596

Fresno, V., Parkes, S. L., Faugère, A., Coutureau, E., & Wolff, M. (2019). A thalamocortical circuit for updating action-outcome associations. ELife, 8, 1–13. https://doi.org/10.7554/eLife.46187

Gerfen, C. R., & Clavier, R. M. (1979). Neural inputs to the prefrontal agranular insular cortex in the rat: Horseradish peroxidase study. Brain Research Bulletin. https://doi.org/10.1016/S0361-9230(79)80012-X

Groenewegen, H. J. (1988). Organization of the afferent connections of the mediodorsal thalamic nucleus in the rat, related to the mediodorsal-prefrontal topography. Neuroscience, 24(2), 379–431. https://doi.org/10.1016/0306-4522(88)90339-9

Hoover, W. B., & Vertes, R. P. (2011). Projections of the medial orbital and ventral orbital cortex in the rat. Journal of Comparative Neurology, 519(18), 3766–3801. https://doi.org/10.1002/cne.22733

Izquierdo, A. (2017). Functional heterogeneity within rat orbitofrontal cortex in reward learning and decision making. Journal of Neuroscience, 37(44), 10529–10540. https://doi.org/10.1523/JNEUROSCI.1678-17.2017

Izquierdo, A., Darling, C., Manos, N., Pozos, H., Kim, C., Ostrander, S., Cazares, V., Stepp, H., & Rudebeck, P. H. (2013). Basolateral amygdala lesions facilitate reward choices after negative feedback in rats. Journal of Neuroscience, 33(9), 4105–4109. https://doi.org/10.1523/JNEUROSCI.4942-12.2013

Johnson, A. W., Gallagher, M., & Holland, P. C. (2009). The basolateral amygdala is critical to the expression of pavlovian and instrumental outcome-specific reinforcer devaluation effects. Journal of Neuroscience. https://doi.org/10.1523/JNEUROSCI.3758-08.2009

Kita, H., & Kitai, S. T. (1990). Amygdaloid projections to the frontal cortex and the striatum in the rat. Journal of Comparative Neurology, 298(1), 40–49. https://doi.org/10.1002/cne.902980104

Krettek, J. E., & Price, J. L. (1977a). Projections from the amygdaloid complex to the cerebral cortex and thalamus in the rat and cat. Journal of Comparative Neurology, 172(4), 687–722. https://doi.org/10.1002/cne.901720408

Krettek, J. E., & Price, J. L. (1977b). The cortical projections of the mediodorsal nucleus and adjacent thalamic nuclei in the rat. Journal of Comparative Neurology. https://doi.org/10.1002/cne.901710204

Kuramoto, E., Iwai, H., Yamanaka, A., Ohno, S., Seki, H., Tanaka, Y. R., Furuta, T., Hioki, H., & Goto, T. (2017). Dorsal and ventral parts of thalamic nucleus submedius project to different areas of rat orbitofrontal cortex: A single neuron-tracing study using virus vectors. Journal of Comparative Neurology, 525(18), 3821–3839. https://doi.org/10.1002/cne.24306

Laubach, M., Amarante, L. M., Swanson, K., & White, S. R. (2018). What, if anything, is rodent prefrontal cortex? ENeuro, 5(5). https://doi.org/10.1523/ENEURO.0315-18.2018

Lenth, R., Singmann, H., Love, J., Buerkner, P., & Herve, M. (2020). Estimated marginal means, aka least-squares means. In R package version 1.5.0.

Lichtenberg, N. T., Pennington, Z. T., Holley, S. M., Greenfield, V. Y., Cepeda, C., Levine, M. S., & Wassum, K. M. (2017). Basolateral amygdala to orbitofrontal cortex projections enable cue-triggered reward expectations. Journal of Neuroscience, 37(35). https://doi.org/10.1523/JNEUROSCI.0486-17.2017

Mcdonald, A. J. (1991). Organization of amygdaloid projections to the prefrontal cortex and associated striatum in the rat. Neuroscience, 44(1), 1–14. https://doi.org/10.1016/0306-4522(91)90247-L

Mcdonald, Alexander J. (1998). Cortical pathways to the mammalian amygdala. Progress in Neurobiology, 55(3), 257–332. https://doi.org/10.1016/S0301-0082(98)00003-3

Murphy, M. J. M., & Deutch, A. Y. (2018). Organization of afferents to the orbitofrontal cortex in the rat. Journal of Comparative Neurology, 526(9), 1498–1526. https://doi.org/10.1002/cne.24424

Murray, E. A., Moylan, E. J., Saleem, K. S., Basile, B. M., & Turchi, J. (2015). Specialized areas for value updating and goal selection in the primate orbitofrontal cortex. ELife. https://doi.org/10.7554/eLife.11695

Ongur, D., & Price, J. L. (2000). The Organization of Networks within the Orbital and Medial Prefrontal Cortex of Rats, Monkeys and Humans. Cerebral Cortex, 10(3), 206–219. https://doi.org/10.1093/cercor/10.3.206

Ostlund, S. B., & Balleine, B. W. (2005). Lesions of medial prefrontal cortex disrupt the acquisition but not the expression of goal-directed learning. Journal of Neuroscience, 25(34), 7763–7770. https://doi.org/10.1523/JNEUROSCI.1921-05.2005

Ostlund, S. B., & Balleine, B. W. (2007). Orbitofrontal cortex mediates outcome encoding in pavlovian but not instrumental conditioning. Journal of Neuroscience, 27(18), 4819–4825. https://doi.org/10.1523/JNEUROSCI.5443-06.2007

Panayi, M. C., & Killcross, S. (2018). Functional heterogeneity within the rodent lateral orbitofrontal cortex dissociates outcome devaluation and reversal learning deficits. ELife, 7, 1–27. https://doi.org/10.7554/eLife.37357.001

Passingham, R. E., Stephan, K. E., & Kötter, R. (2002). The anatomical basis of functional localization in the cortex. Nature Reviews Neuroscience, 3(8), 606–616. https://doi.org/10.1038/nrn893

Paxinos, G., & Watson, C. (1998). The Rat Brain in Stereotaxic Coordinates. In Elsevier Academic Press (4th ed.).

Paxinos, G., & Watson, C. (2013). The Rat Brain in Stereotaxic Coordinates Seventh Edition. Elsevier Academic Press.

Price, J. L. (2006). Architectonic structure of the orbital and medial prefrontal cortex. In The Orbitofrontal Cortex. https://doi.org/10.1093/acprof:oso/9780198565741.003.0001

Price, J. L. (2007). Definition of the Orbital Cortex in Relation to Specific Connections with Limbic and Visceral Structures and Other Cortical Regions. Annals of the New York Academy of Sciences, 1121(1), 54–71. https://doi.org/10.1196/annals.1401.008

R Core Team (2020). (2020). R: A language and environment for statistical computing. In R: A language and environment for statistical computing. R Foundation for Statistical Computing, Vienna, Austria.

Ray, J. P., & Price, J. L. (1992). The organization of the thalamocortical connections of the mediodorsal thalamic nucleus in the rat, related to the ventral forebrain–prefrontal cortex topography. Journal of Comparative Neurology, 323(2), 167–197. https://doi.org/10.1002/cne.903230204

Reep, R. L., Corwin, J. V., & King, V. (1996). Neuronal connections of orbital cortex in rats: Topography of cortical and thalamic afferents. Experimental Brain Research. https://doi.org/10.1007/BF00227299

Rose, J. E., & Woolsey, C. N. (1948). The orbitofrontal cortex and its connections with the mediodorsal nucleus in rabbit, sheep and cat. Research Publications - Association for Research in Nervous and Mental Disease.

Schilman, E. A., Uylings, H. B. M., Graaf, Y. G. de, Joel, D., & Groenewegen, H. J. (2008). The orbital cortex in rats topographically projects to central parts of the caudateputamen complex. Neuroscience Letters, 432(1), 40–45. https://doi.org/10.1016/j.neulet.2007.12.024

Schoenbaum, G., Setlow, B., Saddoris, M. P., & Gallagher, M. (2003). Encoding predicted outcome and acquired value in orbitofrontal cortex during cue sampling depends upon input from basolateral amygdala. Neuron, 39(5), 855–867. https://doi.org/10.1016/S0896-6273(03)00474-4

Sharpe, M. J., & Schoenbaum, G. (2016). Back to basics: Making predictions in the orbitofrontal-amygdala circuit. In Neurobiology of Learning and Memory (Vol. 131). https://doi.org/10.1016/j.nlm.2016.04.009

Stalnaker, T. A., Franz, T. M., Singh, T., & Schoenbaum, G. (2007). Basolateral Amygdala Lesions Abolish Orbitofrontal-Dependent Reversal Impairments. Neuron, 54(1), 51–58. https://doi.org/10.1016/j.neuron.2007.02.014

Takagishi, M., & Chiba, T. (1991). Efferent projections of the infralimbic (area 25) region of the medial prefrontal cortex in the rat: an anterograde tracer PHA-L study. Brain Research, 566(1–2), 26–39. https://doi.org/10.1016/0006-8993(91)91677-S

Tang, J. S., Qu, C. L., & Huo, F. Q. (2009). The thalamic nucleus submedius and ventrolateral orbital cortex are involved in nociceptive modulation: A novel pain modulation pathway. In Progress in Neurobiology. https://doi.org/10.1016/j.pneurobio.2009.10.002

Uylings, H. B. M., Groenewegen, H. J., & Kolb, B. (2003). Do rats have a prefrontal cortex? Behavioural Brain Research, 146(1–2), 3–17. https://doi.org/10.1016/j.bbr.2003.09.028

Van De Werd, H. J. J. M., & Uylings, H. B. M. (2008). The rat orbital and agranular insular prefrontal cortical areas: A cytoarchitectonic and chemoarchitectonic study. Brain Structure and Function, 212(5), 387–401. https://doi.org/10.1007/s00429-007-0164-y

Vertes, R. P. (2004). Differential Projections of the Infralimbic and Prelimbic Cortex in the Rat. Synapse, 51(1), 32–58. https://doi.org/10.1002/syn.10279

Vertes, R. P., & Hoover, W. B. (2008). Projections of the paraventricular and paratenial nuclei of the dorsal midline thalamus in the rat. Journal of Comparative Neurology, 508(2), 212–237. https://doi.org/10.1002/cne.21679

Wobbrock, J. O., Findlater, L., Gergle, D., & Higgins, J. J. (2011). The Aligned Rank Transform for nonparametric factorial analyses using only ANOVA procedures. Conference on Human Factors in Computing Systems - Proceedings. https://doi.org/10.1145/1978942.1978963

Yoshida, A., Dostrovsky, J. O., & Chiang, C. Y. (1992). The afferent and efferent connections of the nucleus submedius in the rat. Journal of Comparative Neurology. https://doi.org/10.1002/cne.903240109

