## Supplemental Information for "Organisation of afferents along the anterior-posterior and medial-lateral axes of the rat orbitofrontal cortex"

### Supplementary materials

#### Supplementary figures

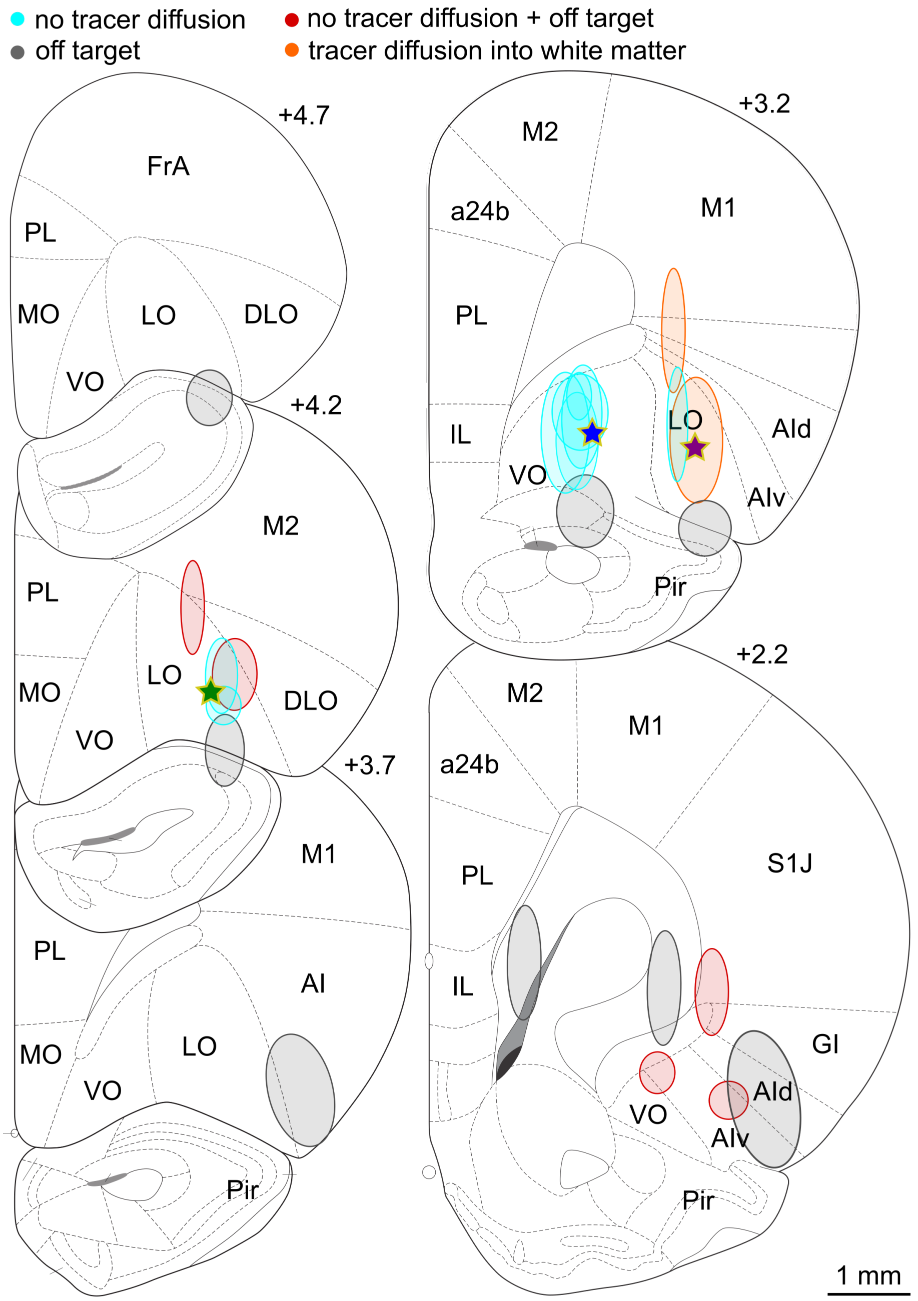


***Figure S1.*** **Coronal sections depicting core of tracer deposits of excluded injections, illustrated at the level of maximal tracer deposit.** Star and oval shapes represent the intended and the observed injection sites core, respectively. Data from injections that were off target ($n$ = 11), in which tracer did not diffuse ($n$ = 8), were both off target and the tracer did not diffuse ($n$ = 6) or diffused into the white matter ($n$ = 2) were excluded from all analyses. Two of the excluded injections were anterior to +4.7 mm from bregma and are not represented here. Distances shown are distances from bregma in mm. a24b: anterior cingulate cortex, area 24b; AI: agranular insular cortex; AId: agranular insular cortex, dorsal part; AIv: agranular insular cortex, ventral part; DLO: orbitofrontal cortex, dorsolateral part;

FrA: frontal association cortex; GI: granular insular cortex; IL: infralimbic cortex; LO: orbitofrontal cortex, lateral part; M1: primary motor cortex; M2: secondary motor cortex; MO: orbitofrontal cortex, medial part; Pir: piriform cortex; PL: prelimbic cortex; VO: orbitofrontal cortex, ventral part; S1J: primary somatosensory cortex, jaw region.

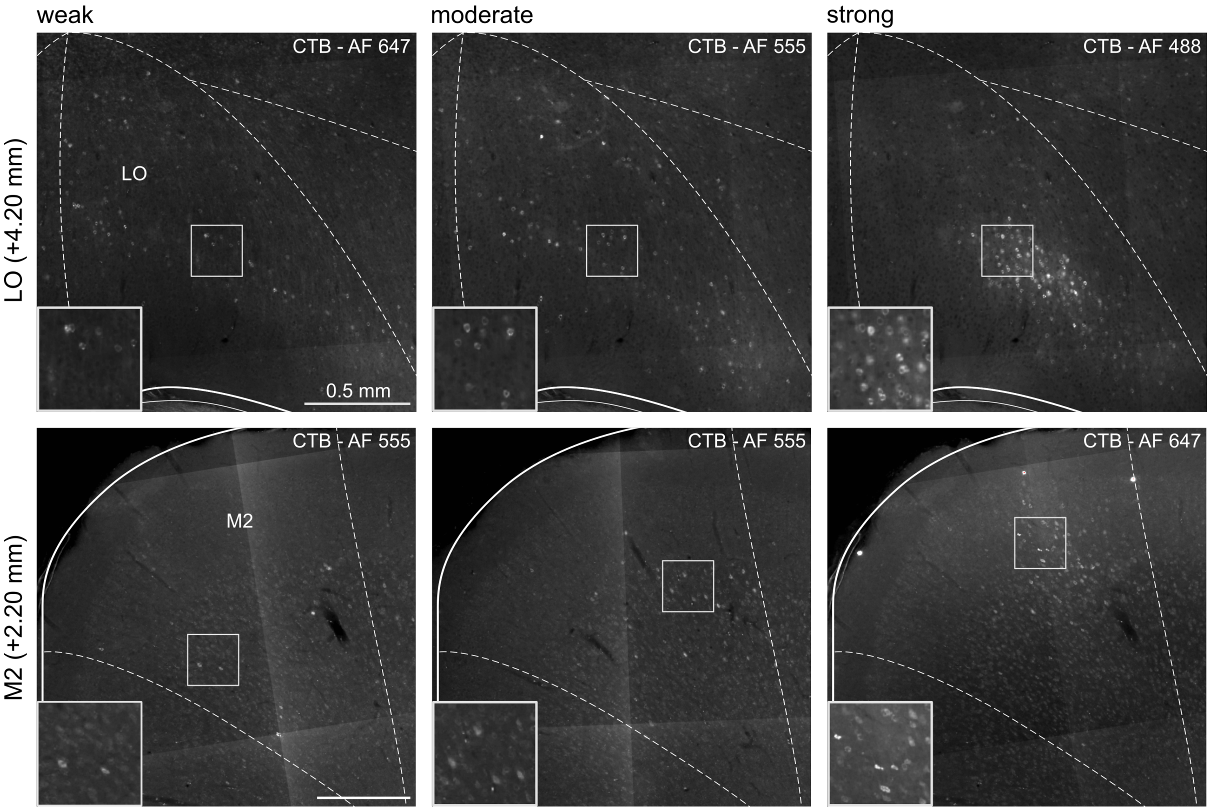


***Figure S2.*** **Example labelling strength classifications.** Depicted are example coronal sections of different brain regions (LO, M2), which labelling strength was classified as weak, moderate, or strong after injection of the retrograde tracer CTB into OFC. Distances shown are distances from bregma in mm. AF: Alexa Fluor; CTB: cholera toxin subunit B; LO: lateral orbitofrontal cortex; M2: secondary motor cortex; OFC: orbitofrontal cortex.

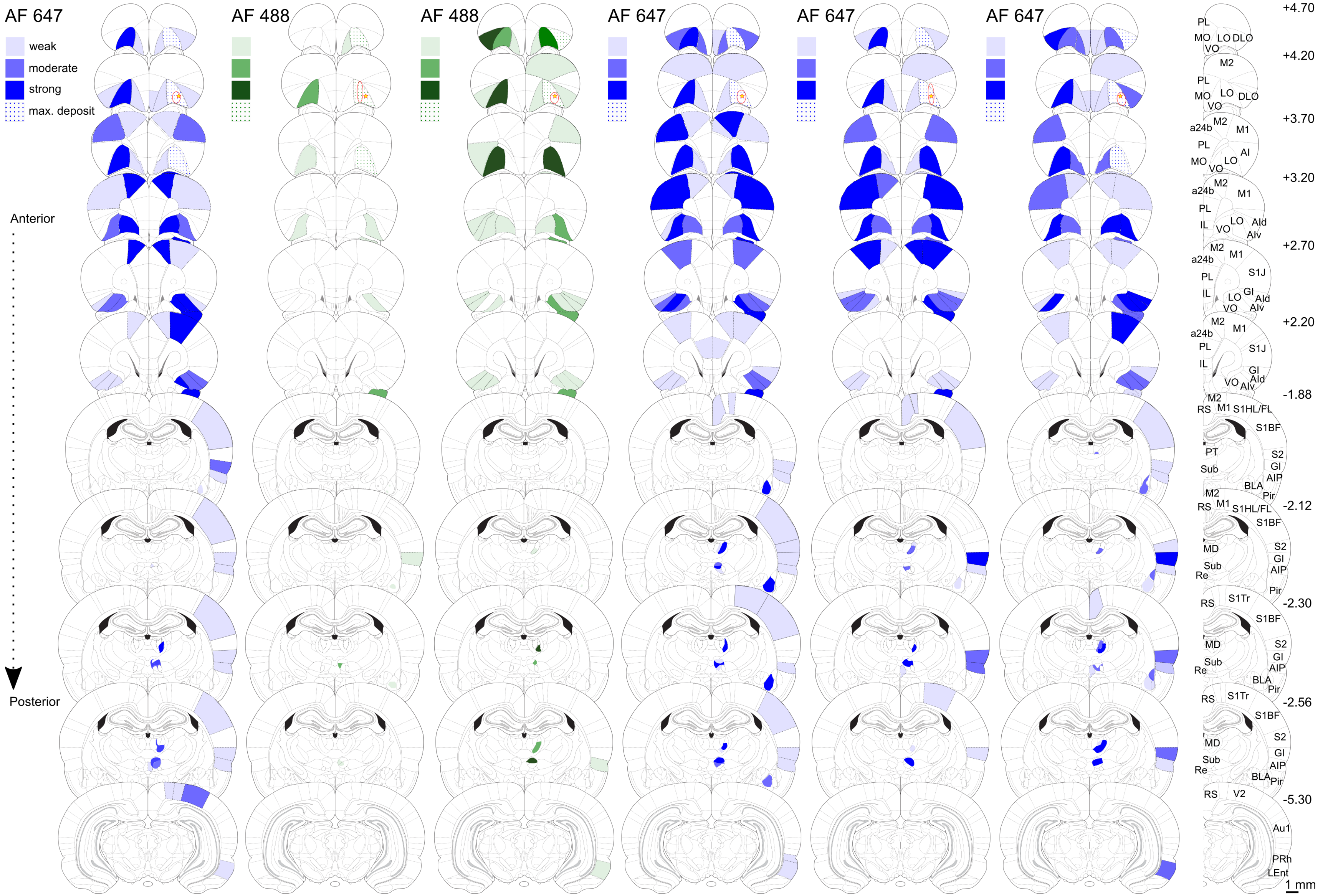
AF 647
weak
moderate
strong
max. deposit
AF 488
AF 488
AF 647
AF 647
AF 647
Anterior
Posterior
+4.70
+4.20
+3.70
+3.20
+2.70
+2.20
-1.88
-2.12
-2.30
-2.56
-5.30
PL
MO
LO
DLO
VO
M2
a24b
M1
AI
IL
AId
AIv
S1J
GI
RS
S1HL/FL
S1BF
PT
Sub
S2
AIP
BLA
Pir
MD
Re
S1Tr
V2
Au1
PRh
LEnt
1 mm

***Figure S3.*** **Density of retrogradely-labelled cells following CTB injection into ALO.** Lighter or darker shades represent lower or higher labelling densities, respectively. Red star and oval shapes represent the intended and the observed injection sites core, respectively. Brains are sorted by the location of its observed injection site, from medial to lateral. Distances shown are distances from bregma in mm. AF: Alexa Fluor; CTB: cholera toxin subunit B; a24b: anterior cingulate cortex, area 24b; ALO: lateral orbitofrontal cortex, anterior part; AI, v, d, P: agranular insular cortex, ventral area, dorsal area, posterior area; Au1: primary auditory cortex; BLA: basolateral amygdala; DLO: dorsolateral orbitofrontal cortex; GI: granular insular cortex; IL: infralimbic cortex; LEnt: lateral entorhinal cortex; LO: lateral orbitofrontal cortex; M1: primary motor cortex; M2: secondary motor cortex; MD: mediodorsal nucleus of the thalamus; MO: medial orbitofrontal cortex; Pir: piriform cortex; PL: prelimbic cortex; PRh: perirhinal cortex; PT: paratenial nucleus of the thalamus; Re: nucleus reuniens of the thalamus; RS: retrosplenial cortex; S1BF: primary somatosensory cortex, barrel field; S1J: primary somatosensory cortex, jaw region; S1HL/FL: primary somatosensory cortex, hindlimb and forelimb regions; S1Tr: primary somatosensory cortex, trunk region; S2: secondary somatosensory cortex; Sub: submedius nucleus of the thalamus; V2: secondary visual cortex; VO: ventral orbitofrontal cortex.

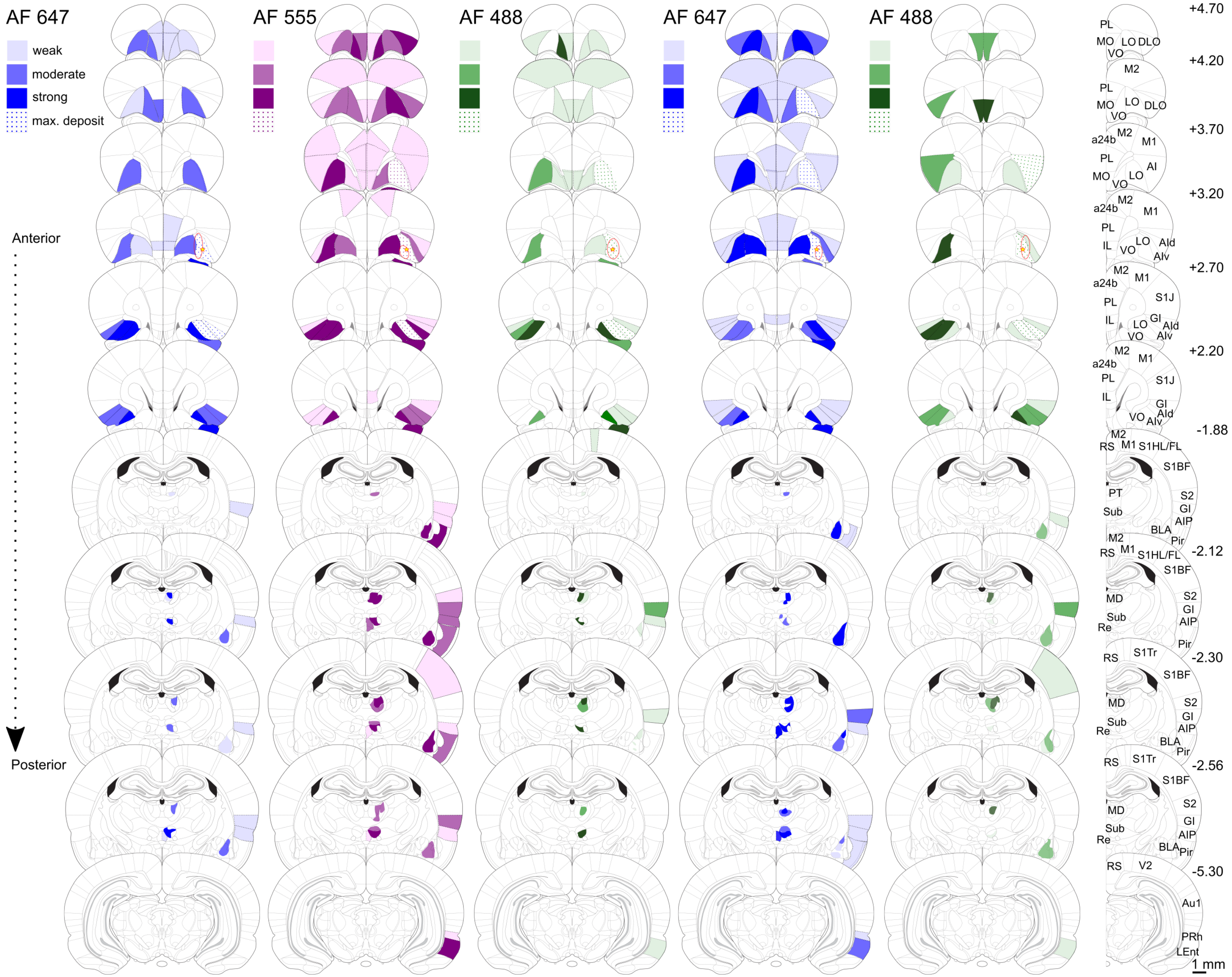
AF 647
weak
moderate
strong
max. deposit
AF 555
AF 488
AF 647
AF 488
Anterior
Posterior
+4.70
+4.20
+3.70
+3.20
+2.70
+2.20
-1.88
-2.12
-2.30
-2.56
-5.30
PL
MO
LO
DLO
VO
M2
a24b
M1
AI
IL
AId
AIv
S1J
GI
RS
S1HL/FL
S1BF
PT
Sub
S2
AIP
BLA
Pir
MD
Re
S1Tr
V2
Au1
PRh
LEnt
1 mm

***Figure S4.*** **Density of retrogradely-labelled cells following CTB injection into PLO.** Lighter or darker shades represent lower or higher labelling densities, respectively. Red star and oval shapes represent the intended and the observed injection sites core, respectively. Brains are sorted by the location of its observed injection site, from medial to lateral. Distances shown are distances from bregma in mm. AF: Alexa Fluor; CTB: cholera toxin subunit B; a24b: anterior cingulate cortex, area 24b; AI, v, d, P: agranular insular cortex, ventral area, dorsal area, posterior area; Au1: primary auditory cortex; BLA: basolateral amygdala; DLO: dorsolateral orbitofrontal cortex; GI: granular insular cortex; IL: infralimbic cortex; LEnt: lateral entorhinal cortex; LO: lateral orbitofrontal cortex; M1: primary motor cortex; M2: secondary motor cortex; MD: mediodorsal nucleus of the thalamus; MO: medial orbitofrontal cortex; Pir: piriform cortex; PL: prelimbic cortex; PLO: lateral orbitofrontal cortex, posterior part; PRh: perirhinal cortex; PT: paratenial nucleus of the thalamus; Re: nucleus reuniens of the thalamus; RS: retrosplenial cortex; S1BF: primary somatosensory cortex, barrel field; S1J: primary somatosensory cortex, jaw region; S1HL/FL: primary somatosensory cortex, hindlimb and forelimb regions; S1Tr: primary somatosensory cortex, trunk region; S2: secondary somatosensory cortex; Sub: submedius nucleus of the thalamus; V2: secondary visual cortex; VO: ventral orbitofrontal cortex.

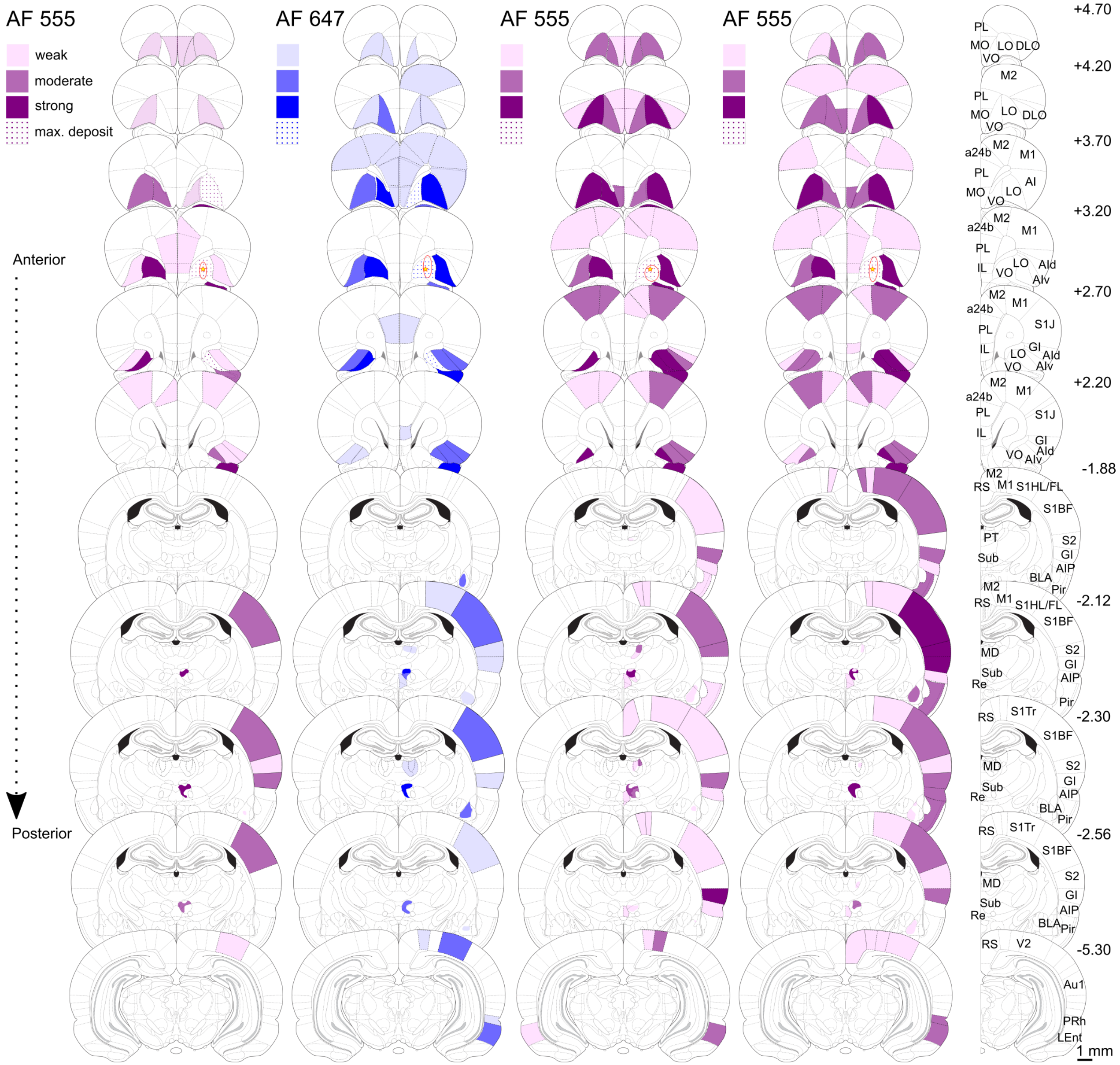
AF 555
weak
moderate
strong
max. deposit
AF 647
AF 555
AF 555
Anterior
Posterior
+4.70
+4.20
+3.70
+3.20
+2.70
+2.20
-1.88
-2.12
-2.30
-2.56
-5.30
PL
MO
LO
DLO
VO
M2
a24b
M1
AI
IL
AId
AIv
S1J
GI
RS
S1HL/FL
S1BF
PT
Sub
S2
AIP
BLA
Pir
MD
Re
S1Tr
V2
Au1
PRh
LEnt
1 mm

***Figure S5.*** **Density of retrogradely-labelled cells following CTB injection into PVO.** Lighter or darker shades represent lower or higher labelling densities, respectively. Red star and oval shapes represent the intended and the observed injection sites core, respectively. Brains are sorted by the location of its observed injection site, from medial to lateral. Distances shown are distances from bregma in mm. AF: Alexa Fluor; CTB: cholera toxin subunit B; a24b: anterior cingulate cortex, area 24b; AI, v, d, P: agranular insular cortex, ventral area, dorsal area, posterior area; Au1: primary auditory cortex; BLA: basolateral amygdala; DLO: dorsolateral orbitofrontal cortex; GI: granular insular cortex; IL: infralimbic cortex; LEnt: lateral entorhinal cortex; LO: lateral orbitofrontal cortex; M1: primary motor cortex; M2: secondary motor cortex; MD: mediodorsal nucleus of the thalamus; MO: medial orbitofrontal cortex; Pir: piriform cortex; PL: prelimbic cortex; PVO: ventral orbitofrontal cortex, posterior part; PRh: perirhinal cortex; PT: paratenial nucleus of the thalamus; Re: nucleus reuniens of the thalamus; RS: retrosplenial cortex; S1BF: primary somatosensory cortex, barrel field; S1J: primary somatosensory cortex, jaw region; S1HL/FL: primary somatosensory cortex, hindlimb and forelimb regions; S1Tr: primary somatosensory cortex, trunk region; S2: secondary somatosensory cortex; Sub: submedius nucleus of the thalamus; V2: secondary visual cortex; VO: ventral orbitofrontal cortex.

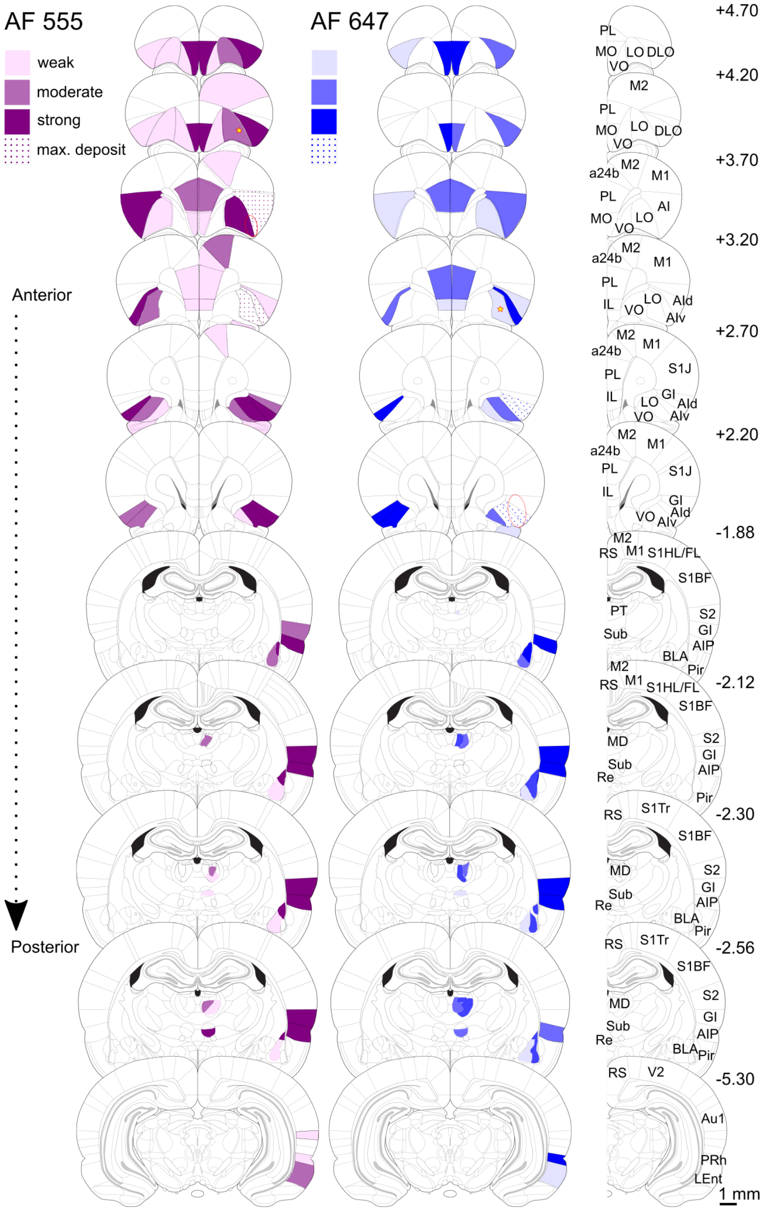


***Figure S6.*** **Density of retrogradely-labelled cells following injection of CTB into AI.** Represented are the labelling patterns of two injections in which the core of the injection extended into AI. Lighter or darker shades represent lower or higher labelling densities, respectively. Red star and oval shapes represent the intended and the observed injection sites core, respectively. Distances shown are distances from bregma in mm. AF: Alexa Fluor; CTB: cholera toxin subunit B; a24b: anterior cingulate cortex, area 24b; AI, v, d, P: agranular insular cortex, ventral area, dorsal area, posterior area; Au1: primary auditory cortex; BLA: basolateral amygdala; DLO: dorsolateral orbitofrontal cortex; GI: granular insular cortex; IL: infralimbic cortex; LEnt: lateral entorhinal cortex; LO: lateral orbitofrontal cortex; M1: primary motor cortex; M2: secondary motor cortex; MD: mediodorsal nucleus of the thalamus; MO: medial orbitofrontal cortex; Pir: piriform cortex; PL: prelimbic cortex; PRh: perirhinal cortex; PT: paratenial nucleus of the thalamus; Re: nucleus reuniens of the thalamus; RS: retrosplenial cortex; S1BF: primary somatosensory cortex, barrel field; S1J: primary somatosensory cortex, jaw region; S1HL/FL: primary somatosensory cortex, hindlimb and forelimb regions; S1Tr: primary somatosensory cortex, trunk region; S2: secondary somatosensory cortex; Sub: submedius nucleus of the thalamus; V2: secondary visual cortex; VO: ventral orbitofrontal cortex.

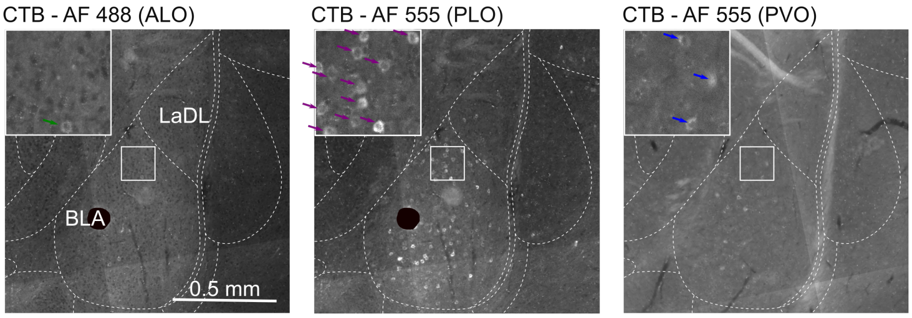


***Figure S7.*** **Labelling in the amygdala following CTB injection into ALO, PLO, or PVO.** Micrographs from representative labelled brains, at -2.12 mm from bregma. We have masked with a black circle a bright artifact present in the slices representing labelling after ALO and PLO injections. ALO: lateral orbitofrontal cortex, anterior area; CTB: cholera toxin subunit B; BLA: basolateral amygdala, anterior part; LaDL: lateral amygdala, dorsolateral part; PLO: lateral orbitofrontal cortex, posterior area; PVO: ventral orbitofrontal cortex, posterior area.

### Supplementary tables

Table S1

*Average labelling density observed following CTB injection into ALO, PLO, or PVO*

| | | Ipsilateral | | | Contralateral | | |
|---|---|---|---|---|---|---|---|
| Structure | AP coordinate (mm) | ALO | PLO | PVO | ALO | PLO | PVO |
| **Prefrontal cortex** | | | | | | | |
| Orbitofrontal cortex | | | | | | | |
| dorsolateral part | +4.70 | 1.00 | 1.00 | 0 | 1.50 | 0.80 | 0 |
| | +4.20 | 0.67 | 0.60 | 0.25 | 0.33 | 0.80 | 0.25 |
| lateral part | +4.70 | m. d. | 1.40 | 1.25 | 2.17 | 1.40 | 1.25 |
| | +4.20 | m. d. | 1.25 | 2.00 | 2.83 | 2.20 | 1.75 |
| | +3.70 | 3.00 | 1.25 | 3.00 | 2.67 | 2.80 | 2.50 |
| | +3.20 | 2.33 | m. d. | 2.50 | 2.33 | 2.80 | 1.75 |
| | +2.70 | 2.00 | m. d. | 2.25 | 1.67 | 1.40 | 1.50 |
| medial part | +4.70 | 0.33 | 1.20 | 0.50 | 0.17 | 1.60 | 0.50 |
| | +4.20 | 0.33 | 1.20 | 0.50 | 0.33 | 1.40 | 0.75 |
| | +3.70 | 0 | 1.00 | 0.75 | 0 | 0.80 | 0.75 |
| ventral part | +4.70 | 1.00 | 1.40 | 1.50 | 1.00 | 2.00 | 1.25 |
| | +4.20 | 0.83 | 1.00 | 1.25 | 0.67 | 1.40 | 1.50 |
| | +3.70 | 0.83 | 0.80 | 1.67 | 0.83 | 0.60 | 2.50 |
| | +3.20 | 1.83 | 1.80 | m. d. | 1.67 | 1.60 | 3.00 |
| | +2.70 | 2.17 | 2.40 | 3.00 | 1.00 | 1.80 | 2.75 |
| | +2.20 | 0 | 2.80 | 2.50 | 0 | 1.80 | 1.50 |
| Infralimbic cortex | +3.20 | 0 | 0.40 | 0.50 | 0 | 0.40 | 0.50 |
| | +2.70 | 0 | 0.20 | 0.25 | 0 | 0.20 | 0 |
| | +2.20 | 0 | 0.20 | 0.25 | 0 | 0 | 0 |
| Prelimbic cortex | +4.70 | 0 | 0.20 | 0 | 0 | 0.40 | 0 |
| | +4.20 | 0 | 0.20 | 0.25 | 0 | 0.20 | 0.25 |
| | +3.70 | 0 | 0.40 | 0.25 | 0 | 0.40 | 0.25 |
| | +3.20 | 0 | 0.40 | 0.50 | 0 | 0.20 | 0.25 |
| | +2.70 | 0.17 | 0 | 0.25 | 0.17 | 0 | 0.25 |
| | +2.20 | 0 | 0 | 0 | 0 | 0 | 0.25 |
| Cingulate cortex, area 1 | +3.70 | 0.17 | 0.20 | 0.50 | 0 | 0.20 | 0 |
| | +3.20 | 0 | 0 | 0.75 | 0 | 0 | 0 |
| | +2.70 | 0.17 | 0 | 0.25 | 0 | 0 | 0.25 |
| | +2.20 | 0 | 0 | 0.25 | 0 | 0 | 0 |
| **Insular cortex** | | | | | | | |
| Agranular insular cortex | +3.70 | 0 | 0.50 | 0.20 | 0.17 | 0.80 | 0 |
| | -1.88 | 0.67 | 0.40 | 0.50 | 0 | 0 | 0 |
| | -2.12 | 0.83 | 0.40 | 0.25 | 0 | 0 | 0 |
| | -2.30 | 0.67 | 0.60 | 0.75 | 0 | 0 | 0 |
| | -2.56 | 0.50 | 0.60 | 0.25 | 0 | 0 | 0 |
| dorsal part | +3.20 | 0 | 0.40 | 0 | 0.17 | 0.20 | 0 |
| | +2.70 | 1.17 | 0.80 | 0.25 | 0.67 | 1.20 | 0 |
| | +2.20 | 1.33 | 1.80 | 1.75 | 0.83 | 1.00 | 0 |
| ventral part | +3.20 | 0.17 | 1.00 | 0 | 0.33 | 0.40 | 0 |
| | +2.70 | 1.50 | m. d. | 1.25 | 1.17 | 2.20 | 0.25 |
| | +2.20 | 1.33 | 1.80 | 1.75 | 0.83 | 1.20 | 0.50 |
| Granular insular cortex | +2.70 | 0.17 | 0.40 | 0 | 0.17 | 0.20 | 0 |
| | +2.20 | 0.33 | 0.60 | 0 | 0 | 0.20 | 0 |
| | -1.88 | 0.83 | 0.40 | 0.50 | 0 | 0 | 0 |
| | -2.12 | 0.17 | 1.20 | 1.50 | 0 | 0 | 0 |
| | -2.30 | 0.83 | 1.00 | 1.75 | 0 | 0 | 0 |
| | -2.56 | 0.83 | 0.80 | 1.25 | 0 | 0 | 0 |
| **Motor cortex** | | | | | | | |
| Primary | +3.70 | 1.00 | 0.20 | 0.50 | 1.67 | 0.20 | 0.50 |
| | +3.20 | 1.33 | 0 | 0.50 | 1.50 | 0 | 1.00 |
| | +2.70 | 1.17 | 0 | 1.00 | 1.00 | 0 | 1.00 |
| | +2.20 | 1.17 | 0 | 1.00 | 0.33 | 0 | 0 |
| | -1.88 | 0.17 | 0.20 | 0.25 | 0 | 0 | 0 |
| | -2.12 | 0 | 0 | 0.50 | 0 | 0 | 0 |
| | -2.30 | 0 | 0 | 0 | 0 | 0 | 0 |
| | -2.56 | 0 | 0 | 0.25 | 0 | 0 | 0 |
| Secondary | +4.20 | 0.67 | 0.60 | 0.50 | 0.17 | 0.60 | 0.25 |
| | +3.70 | 0.67 | 0.20 | 0.25 | 0.33 | 0 | 0.25 |
| | +3.20 | 1.00 | 0.20 | 0.25 | 1.33 | 0.20 | 0.50 |
| | +2.70 | 1.33 | 0.20 | 0.50 | 0.83 | 0 | 1.00 |
| | +2.20 | 0.33 | 0 | 0.75 | 0.17 | 0 | 0.50 |
| | -1.88 | 1.00 | 0 | 0.50 | 0 | 0 | 0.25 |
| | -2.12 | 0 | 0 | 2.00 | 0 | 0 | 1.00 |
| | -2.30 | 0 | 0 | 0.25 | 0 | 0 | 0 |
| | -2.56 | 0 | 0 | 0 | 0 | 0 | 0 |
| **Sensory cortices** | | | | | | | |
| Piriform cortex | +3.70 | 1.50 | 1.20 | 2.75 | 0 | 0 | 0 |
| | +3.20 | 2.17 | 1.80 | 2.75 | 0 | 0 | 0 |
| | +2.70 | 2.17 | 2.00 | 3.00 | 0.14 | 0 | 0 |

| | | | | | | | |
|---|---|---|---|---|---|---|---|
| | +2.20 | 2.33 | 2.20 | 0 | 0 | 0 | 0 |
| | -1.88 | 0.17 | 0.80 | 0.75 | 0 | 0 | |
| | -2.12 | 0 | 0.40 | 0.75 | 0 | 0 | 0 |
| | -2.30 | 0 | 0.40 | 0.50 | 0 | 0 | 0 |
| | -2.56 | 0.40 | 0.25 | 0 | 0 | 0 | 0 |
| Somatosensory cortex | | | | | | | |
| primary/barrel cortex | -1.88 | 0.50 | 0 | 0.75 | 0 | 0 | 0 |
| | -2.12 | 0.33 | 0 | 2.25 | 0 | 0 | 0 |
| | -2.30 | 0.33 | 0.40 | 1.75 | 0 | 0 | 0 |
| | -2.56 | 0.33 | 0 | 1.50 | 0 | 0 | 0 |
| forelimb and hindlimb regions | -1.88 | 0 | 0 | 0.50 | 0 | 0 | 0 |
| | -2.12 | 0 | 0 | 0.50 | 0 | 0 | 0 |
| trunk region | -2.30 | 0.17 | 0 | 0.50 | 0 | 0 | 0 |
| | -2.56 | 0.17 | 0 | 0.50 | 0 | 0 | 0 |
| secondary | -1.88 | 0 | 0.20 | 0.25 | 0 | 0 | 0 |
| | -2.12 | 0.40 | 0.20 | 1.50 | 0 | 0 | 0 |
| | -2.30 | 0 | 0 | 0.50 | 0 | 0 | 0 |
| | -2.56 | 0 | 0 | 0.25 | 0 | 0 | 0 |
| Auditory cortex | | | | | | | |
| primary | -5.30 | 0 | 0 | 0 | 0 | 0 | 0 |
| secondary | | | | | | | |
| dorsal part | -5.30 | 0 | 0 | 0 | 0 | 0 | 0 |
| ventral part | -5.30 | 0 | 0 | 0 | 0 | 0 | 0 |
| Visual cortex | | | | | | | |
| primary | | | | | | | |
| binocular area | -5.30 | 0 | 0 | 0 | 0 | 0 | 0 |
| monocular area | -5.30 | 0 | 0 | 1.00 | 0 | 0 | 0 |
| secondary | | | | | | | |
| lateral area | -5.30 | 0.33 | 0 | 0.75 | 0 | 0 | 0 |
| mediolateral area | -5.30 | 0.17 | 0 | 0.75 | 0 | 0 | 0 |
| mediomedial area | -5.30 | 0.17 | 0 | 0.25 | 0 | 0 | 0 |
| **Temporal lobe** | | | | | | | |
| Amygdala | | | | | | | |
| basolateral | -1.88 | 1.17 | 1.60 | 0.75 | 0 | 0 | 0 |
| anterior part | -1.88 | 1.17 | 1.60 | 0.75 | 0 | 0 | 0 |
| | -2.12 | 1.33 | 2.00 | 1.25 | 0 | 0 | 0 |
| | -2.30 | 0.67 | 1.80 | 1.25 | 0 | 0 | 0 |
| | -2.56 | 0.33 | 1.20 | 0.50 | 0 | 0 | 0 |
| posterior part | -2.30 | 0 | 0.40 | 0 | 0 | 0 | 0 |
| | -2.56 | 0 | 0.80 | 0 | 0 | 0 | 0 |
| lateral | | | | | | | |
| dorsolateral part | -1.88 | 0.17 | 1.20 | 0.25 | 0 | 0 | 0 |
| | -2.12 | 0.17 | 1.40 | 0 | 0 | 0 | 0 |
| | -2.30 | 0.50 | 1.20 | 0.25 | 0 | 0 | 0 |
| | -2.56 | 0.17 | 0.40 | 0 | 0 | 0 | 0 |
| ventromedial part | -2.56 | 0 | 0.60 | 0 | 0 | 0 | 0 |
| ventrolateral part | -2.56 | 0 | 0.20 | 0 | 0 | 0 | 0 |
| Entorhinal cortex, lateral part | -5.30 | 0.83 | 1.20 | 1.50 | 0 | 0 | 0.33 |
| Perirhinal cortex | -5.30 | 0.17 | 0.40 | 0.75 | 0 | 0 | 0 |
| **Parietal lobe** | | | | | | | |
| Retrosplenial cortex | -1.88 | 0.33 | 0 | 0 | 0 | 0 | 0 |
| | -2.12 | 0 | 0 | 0 | 0 | 0 | 0 |
| | -2.30 | 0.17 | 0 | 0.25 | 0 | 0 | 0 |
| | -2.56 | 0 | 0 | 0 | 0 | 0 | 0 |
| **Thalamic nuclei** | | | | | | | |
| Mediodorsal | -1.88 | 0 | 0 | 0 | 0 | 0 | 0 |
| central part | -2.30 | 0.33 | 2.20 | 0.75 | 0 | 0 | 0 |
| | -2.56 | 0.83 | 2.60 | 0.25 | 0 | 0 | 0 |
| lateral part | -2.12 | 1.20 | 1.80 | 1.25 | 0 | 0 | 0 |
| | -2.30 | 2.00 | 1.40 | 0.75 | 0 | 0 | 0 |
| | -2.56 | 2.00 | 0.80 | 0 | 0 | 0 | 0 |
| medial part | -2.12 | 0.20 | 2.60 | 0.25 | 0 | 0 | 0 |
| | -2.30 | 0.17 | 1.60 | 0.25 | 0 | 0 | 0 |
| | -2.56 | 0.50 | 0.40 | 0 | 0 | 0 | 0 |
| Paratenial | -1.88 | 0.40 | 1.60 | 0 | 0 | 0 | 0 |
| Reunens | -2.12 | 0 | 0.40 | 0.50 | 0 | 0 | 0 |
| | -2.30 | 0.33 | 0.80 | 0.50 | 0 | 0 | 0 |
| | -2.56 | 0.17 | 0.80 | 0.50 | 0 | 0 | 0 |
| Submedius | -1.88 | 0 | 0 | 0 | 0 | 0 | 0 |
| dorsal part | -2.12 | 0.60 | 0.20 | 2.25 | 0 | 0 | 0 |
| | -2.30 | 1.50 | 0.60 | 3.00 | 0 | 0 | 0 |
| | -2.56 | 2.33 | 1.20 | 1.25 | 0 | 0 | 0 |
| ventral part | -2.12 | 0.60 | 2.40 | 1.25 | 0 | 0 | 0 |
| | -2.30 | 1.50 | 2.60 | 2.50 | 0 | 0 | 0 |
| | -2.56 | 1.00 | 2.40 | 1.25 | 0 | 0 | 0 |

*Note*. Average values were obtained following 4-level categorisation (0, absence of labelling; 1, weak; 2, moderate; 3, strong) of labelling density. Areas with maximal tracer deposit, only found around injection sites, were not included in the analyses. m. d.: maximal deposit.
